# A novel Bayesian model for assessing intratumor heterogeneity of tumor infiltrating leukocytes with multi-region gene expression sequencing

**DOI:** 10.1101/2023.10.24.563820

**Authors:** Peng Yang, Shawna M. Hubert, P. Andrew Futreal, Xingzhi Song, Jianhua Zhang, J. Jack Lee, Ignacio Wistuba, Ying Yuan, Jianjun Zhang, Ziyi Li

## Abstract

Intratumor heterogeneity (ITH) of tumor-infiltrated leukocytes (TILs) is an important phenomenon of cancer biology with potentially profound clinical impacts. Multiregion gene expression sequencing data provide a promising opportunity that allows for explorations of TILs and their intratumor heterogeneity for each subject. Although several existing methods are available to infer the proportions of TILs, considerable methodological gaps exist for evaluating intratumor heterogeneity of TILs with multi-region gene expression data. Here, we develop ICeITH, <u>i</u>mmune <u>c</u>ell <u>e</u>stimation reveals <u>i</u>ntratumor <u>h</u>eterogeneity, a Bayesian hierarchical model that borrows cell type profiles as prior knowledge to decompose mixed bulk data while accounting for the within-subject correlations among tumor samples. ICeITH quantifies intratumor heterogeneity by the variability of targeted cellular compositions. Through extensive simulation studies, we demonstrate that ICeITH is more accurate in measuring relative cellular abundance and evaluating intratumor heterogeneity compared with existing methods. We also assess the ability of ICeITH to stratify patients by their intratumor heterogeneity score and associate the estimations with the survival outcomes. Finally, we apply ICeITH to two multi-region gene expression datasets from lung cancer studies to classify patients into different risk groups according to the ITH estimations of targeted TILs that shape either pro- or anti-tumor processes. In conclusion, ICeITH is a useful tool to evaluate intratumor heterogeneity of TILs from multi-region gene expression data.

## 1 Introduction

Cancers are composed of cells with distinct molecular and phenotypic features within the same tumors, a phenomenon termed intratumor heterogeneity (Zhang et al., 2014). Intratumor heterogeneity (ITH) is a fundamental phenomenon that presents at different molecular levels, between different cancer types, as well as in the constitution of the tumor microenvironment (Sousa et al., 2018). Previous studies have demonstrated that intratumor heterogeneities are associated with an increased risk of post-surgical recurrence in localized non-small cell lung cancers (NSCLS) (Reuben et al., 2017; Gaudreau et al., 2021). Tumor-infiltrated leukocytes (TILs), as a key component of the tumor microenvironment, is a dynamic process that drives intratumor heterogeneity (Jia et al., 2018) and is closely associated with patient prognosis (Sato et al., 2005; Bindea et al., 2013). Tremendous efforts have been put into the research of local mutation diversity (Jia et al., 2018), gene expression variation (Meister et al., 2014) and the evolution of the tumor-microenvironment (Janiszewska et al., 2020). However, none of these studies have systematically studied the intratumor heterogeneities of TILs, which may provide valuable insights into cancer biology and personalized cancer treatment.

Both experimental and computational approaches are available for studying the compositions of TILs. Experimental methods sort the cell to purified samples or rely on single cell technology (Hwang et al., 2018; Potter et al., 2018; Saliba et al., 2014; Bacher et al., 2016; Chen et al., 2019). But these tools are often too expensive and time-consuming in population-scale studies. TIL compositions can also be inferred using computational approaches from the bulk transcriptomic sequencing data (Newman et al., 2015; Rooney et al., 2015; Racle et al., 2017; Li et al., 2016). Most of these methods use a regression-based framework to measure the relative immune cell type abundance with a purified cell-typespecific gene expression as reference. For example, CIBERSORT applies linear support vector regression (SVR) (Schölkopf et al., 2000) using a matrix of reference gene signatures as covariates and estimates the cellular compositions (Newman et al., 2015); EPIC is based on weighted least squares and imposes higher weights on informative genes to improve estimation accuracy (Racle et al., 2020); and TIMER employs a constrained least squares fitting on immune-specific informative genes to remove the potential noises from the tumor cells (Li et al., 2016). However, none of these methods are designed for the problem of estimating intratumor heterogeneity.

Our work is motivated by the problem of estimating intratumor heterogeneity of TILs for lung cancer patients with multi-region RNA sequencing (RNA-seq) data. The two motivating studies are the TRAcking Non-small Cell Lung Cancer Evolution Through Therapy (TRACERx) study (Jamal-Hanjani et al., 2017) and the MD Anderson intratumor heterogeneity (ITH) study for lung cancer (Zhang et al., 2014). Both studies obtained multi-region RNA-seq data from lung cancer patients with each patient providing multiple tumor sample slices. Moreover, TRACERx study provides multi-region matched RNA-seq and WES data in bulk level from 100 patient cohorts diagnosed as early stage non-small cell lung cancers. Current studies focus on investigating the ITH on DNA data only, and they discovered a strong relation in the percentage of subclonal copy number alterations (SCNA) to patient survival, indicating the ITH within each patient plays an important role in cancer progression. However, it is important to note that the methods mentioned earlier were primarily designed to uncover the proportions of TILs by analyzing independent transcriptomic samples. When multiple RNA-seq samples exist per patient, the within-subject correlations are important factors to be accounted for in the estimation of intratumor heterogeneity. Consequently, the existing methods cannot accurately capture the intratumor heterogeneity of TILs in such scenarios. Notably, the ITH of TILs plays an important role in the tumor microenvironment and may be associated with patients’ outcomes. Compared with regular RNA-seq data using one tumor sample per patient, our data have a few unique attributes that the existing methods cannot address. First, the TIL compositions from multiple samples of the same subject tend to be more correlated than those from samples of several different individuals. Such within-subject correlations provide additional information about the estimation of TIL compositions and should be considered in the modeling. Second, the standard regression-based approaches do not consider gene-specific variability and may not be robust. It is likely that such pure cell type profiles have variations across different subjects. Some existing work has recognized this, e.g., BLADE (Andrade et al., 2021) takes a hierarchical Bayesian approach and models gene expression by log-normal distribution to account for the gene-specific variability for each cell type. Still, the method could not address the within-subject correlation in the multi-region data for intratumor heterogeneity evaluation.

In this paper, we develop a novel Bayesian hierarchical model, ICeITH, to infer relative cell type abundance and its variability across bulk tumor samples obtained from a multi-region sequencing design. ICeITH is a reference-based deconvolution method and it overcomes the limitations of current methods by modeling a patient-specific mean expression to account for the heterogeneity of gene expressions introduced from multi-region sequencing design. In addition, ICeITH measures the intratumor heterogeneity by quantifying the variability of targeted cellular composition and it potentially reveals the relation with the risk of patients’ survival. We present the notation, model, and estimations in Section 2. The simulation setups and results for evaluating the empirical performance are described in Section 3. The results of analyzing the two lung cancer multi-region datasets are presented in Section 4. We provide conclusion remarks and discussions in Section 5. The detailed derivations for the Bayesian model are relegated to the Supplementary Material. The proposed methods are available through a user-friendly R package (https://github.come/pengyang0411/ICeITH).

## 2 Method

### 2.1 Input data

ICeITH model is designed to address the challenges of assessing intratumor heterogeneity using multi-region RNA-seq data. Multi-region gene expression data are obtained by sequencing multiple sample slices from the same subject and provide a promising opportunity to explore intratumor heterogeneity, as they can reveal differences in gene expression and immune cell infiltration between different regions of the same tumor. Compared to traditional single-region RNA-seq, multi-region RNA-seq data are generally more affordable. Meanwhile, it allows for a more comprehensive analysis of intratumor heterogeneity than using single-region RNA-seq data. Our motivating studies, TRACERx and MDA-ITH, contain multi-region RNA-seq data from lung cancer patients, which we use to develop and validate ICeITH. Real data evidence from these studies demonstrates a higher correlation for the tumor slices from the same patient by visualizing the similarities of samples within and between different subjects in TRACERx (Figure 1). Compared with other existing methods, ICeITH can borrow information through these higher within-subject correlations and achieve higher accuracy in estimating the patient-specific cell type profiles.

**Figure 1.**
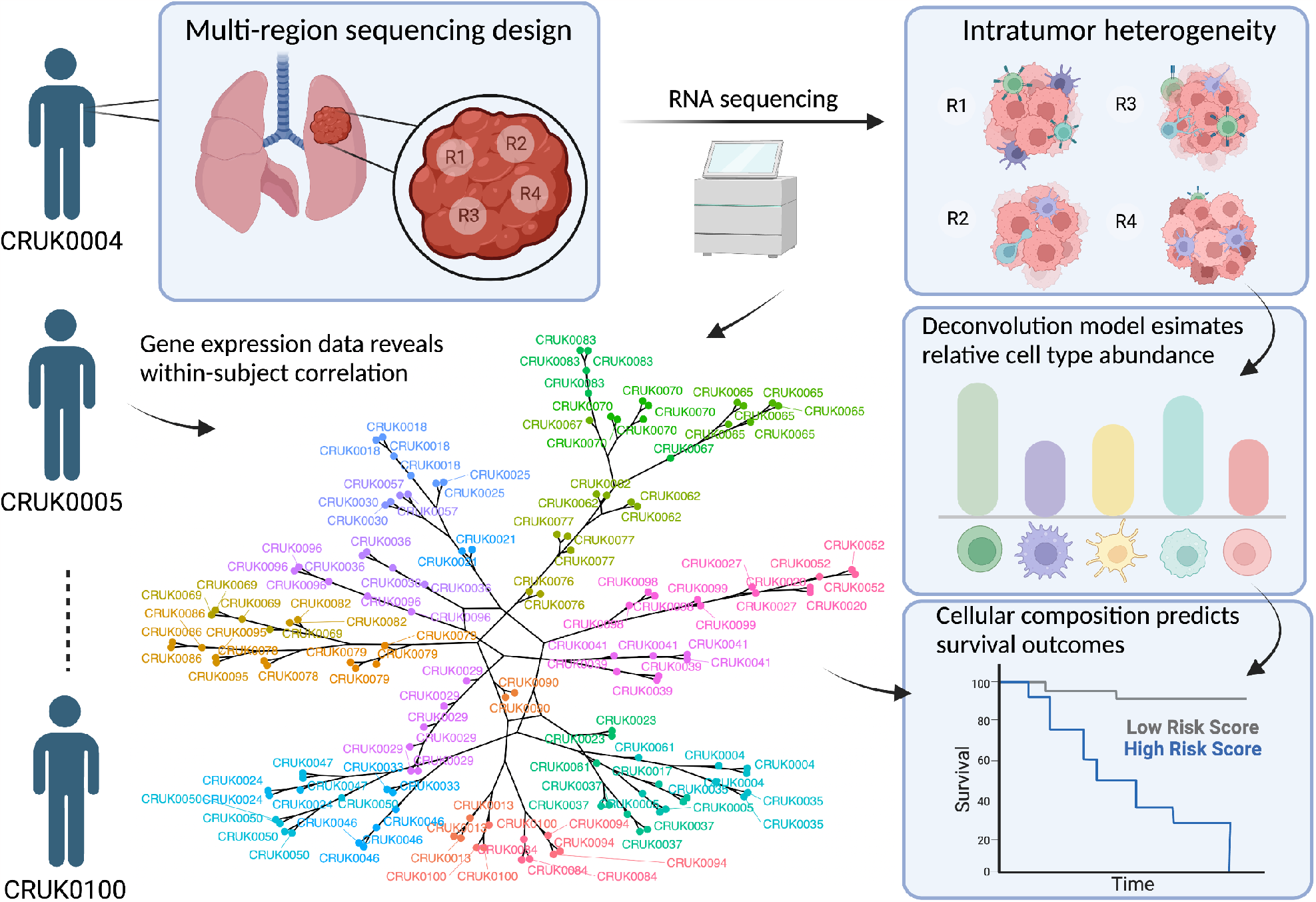
Multi-region sequencing study design. TRACERx data are used to exemplify the data collection procedure and sample relationships. The clustering results by phylogenetic tree show strong within-subject correlations. The proposed deconvolution model borrows information across samples and assesses the intratumor heterogeneity of cellular compositions, which are found to be associated with patients’ prognoses.

In addition to the multi-region gene expression data, ICeITH utilizes purified reference samples to estimate the proportion of immune cell types present in the tumor samples. These reference samples can be obtained from publicly available gene expression data. Several studies have provided immune-specific marker gene sets and their expression profiles (Newman et al., 2015; Racle et al., 2020). Detailed information on constructing major immune cell references using these marker gene sets is provided in Section 4.1. The use of these reference samples and marker gene sets allows ICeITH to accurately estimate the proportions of immune cell types present in the tumor samples and to quantify the intratumor heterogeneity based on these cell types.

### 2.2 Notation

Consider a study with *N* patients. Let *I*_*i*_ denote the sample index for subject *i, i* = 1, 2, …, *N*, and thus subject *i* has |*I*_*i*_|(|*I*_*i*_| *>* 1) tumor slices sequenced by RNA-seq. As tumor samples consist of many cell types including malignant cells, stromal cells, and various immune cells, the gene expression data for each sample can be considered as mixed signals. With additional information, such as genes with immune-specific expressions, we aim to estimate immune cell-type-specific abundance and quantify the intratumor heterogeneity for each patient subject. Several informative immune-specific gene sets are available from previous work, for example, CIBERSORT and EPIC (Newman et al., 2015; Racle et al., 2020). We obtain these gene sets and specifically focus our ICeITH model on the immune-specific genes only. We use ***Y*** and ***X*** to denote the observed gene expressions for mixed bulk tumor samples and purified reference samples for the immune-specific genes, respectively. *Y* is a 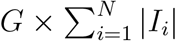 matrix, where *Y*_*sg*_ is the expression of gene *g* in sample *s*. If this sample *s* falls into index *I*_*i*_ (i.e., *s* ∈ *I*_*i*_), it indicates the sample is collected from patient *i*. ***X*** is a three-dimensional tensor of size *V* × *G* × *K*, where *X*_*vgk*_ is the expression of gene *g* in the *v*-th purified reference sample from cell type *k*. The number of genes is denoted by *G*, the number of purified reference samples by *V*, and the number of immune cell types by *K*. An overview of our notations and their association in our model is presented in Figure 2.

**Figure 2.**
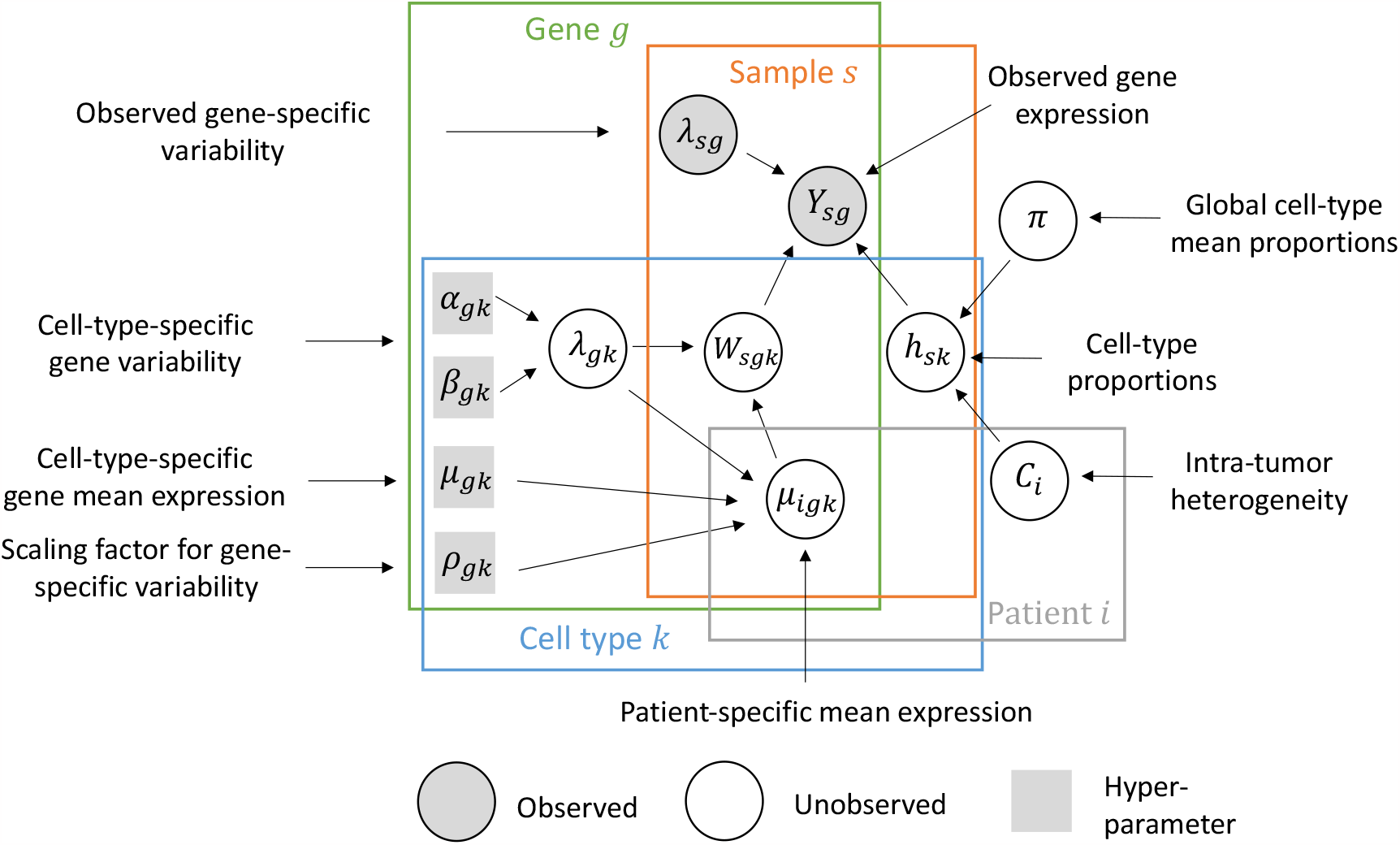
Graphical model structure of ICeITH model.

### 2.3 Model

We begin by modeling the observed cell-type-specific reference *X*_*vgk*_ using a log-normal distribution with mean and precision parameters 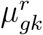 and 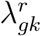, respectively. We obtain these parameters directly from the mean and variance of the log(*X*_*vgk*_) values (Wilson et al., 2020). We then use the estimated 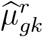 and 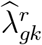 as prior knowledge to guide the deconvolution process.

Considering a total of *K* immune cell types are of interest here, the observed gene expression *Y*_*sg*_ can be decomposed as:

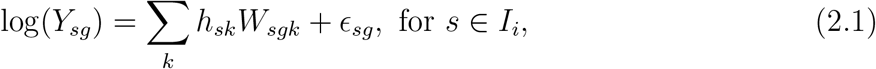

where *h*_*sk*_ is the unobserved cellular abundance from cell type *k* in sample *s*, and *W*_*sgk*_ is a three-dimensional tensor that stands for the hidden expression profiles of gene *g* in sample *s* from cell type *k. ϵ*_*sg*_ is the error term that follows a normal distribution with a mean value of 0 and variance of 1*/ω*_*g*_, where *ω*_*g*_ is estimated empirically by taking the reciprocal of the standard deviation of log(*Y*_*·g*_). The ICeITH model utilizes this weighting scheme to flatten the likelihood of genes, particularly those with larger variance and suffering from over-dispersion, which is a common characteristic of RNA-seq data (Wilson et al., 2020). This approach down-weights the contributions of such genes during the estimation of cell type abundance. This linear assumption frames the ICeITH model as a mixture model problem, where the goal is to estimate the cellular compositions *h*_*sk*_ for each immune cell type *k*.

To characterize intratumor heterogeneity within the same patient subject, a hierarchical Bayesian approach is taken in two steps. First, each patient is allowed to have their own pure cell type profile parameters *μ*_*igk*_ per gene and per cell type. This is expressed as:

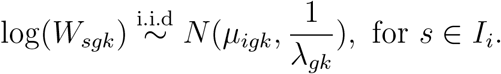

Here, the unobserved gene expression of sample *s* ∈ *I*_*i*_ is assumed to follow a log-normal distribution with its mean centered to its patient-specific mean level, and the discrepancy can be accounted for by *λ*_*gk*_, which is the cell-type-specific variability for each gene. The choice of *λ*_*gk*_ is guided by the reference profiles since the reference samples are collected independently across patients and capture an overall discrepancy in the cell-type-specific genes. Second, we use Dirichlet distribution to model cell type proportion parameters, *h*_*sk*_. For sample *s* ∈ *I*_*i*_, we have

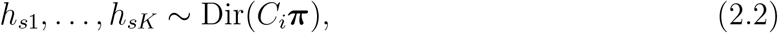

where ***π*** is a *K* by 1 vector pooled across all samples with Σ _*k*_ *π*_*k*_ = 1 to represent the global cellular composition, and *C*_*i*_ is a patient-specific parameter that controls the variability of the cellular composition across samples within each patient, thereby revealing intratumor heterogeneity. When *C*_*i*_ tends to be small, it indicates that patient *i* has greater variability in cellular composition, whereas when *C*_*i*_ increases, it reveals a more homogeneous cellular composition.

The Fenton-Wilkinson (FW) approximation is used to approximate equation (2.1) by another log-normal distribution since there is no closed-form solution for the summation of independent log-normal variables. Specifically, the observed gene expression *Y*_*sg*_ is approximated as follows:

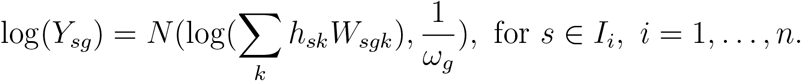

Similar modeling strategies have also been adopted in previous work for deconvolution (Wilson et al., 2020; Andrade et al., 2021).

### 2.4 Prior specifications

To incorporate prior knowledge from existing reference profiles, we use conjugate prior distributions for the patient-specific mean expression parameter *μ*_*igk*_’s and variability *λ*_*gk*_ of each gene *g* and cell type *k*:

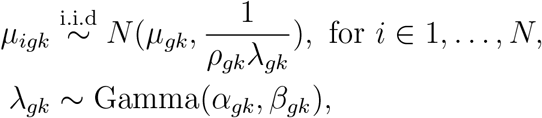

where *ρ*_*gk*_ controls the amount of prior information we borrow from the reference profiles. Larger values of *ρ*_*gk*_ indicate more confidence in prior knowledge and greater borrowing. We estimate the cell-type-specific mean expression parameter *μ*_*gk*_ as the sample mean of the cell-type-specific expression levels in the reference data, denoted by 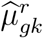. To borrow the cell-type-specific variability from the reference data, we set *α*_*gk*_ = 1 and 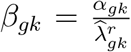, where 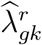 is the sample variance of the cell-type-specific expression levels in the reference data. This choice of hyperparameters ensures that the expected value of *λ*_*gk*_ is equal to 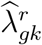.

### 2.5 Optimization

We use the Collapsed Variational Bayesian (CVB) method to optimize the ICeITH model. The CVB method is particularly suitable for models with conjugate priors, as we can collapse the hidden variables and then apply a variational Bayesian inference algorithm. This approach is more computationally efficient than Markov chain Monte Carlo (MCMC). The collapsed space of unobserved variables allows for better satisfaction of mean-field assumptions and results in a more accurate approximation of the true posterior distribution. This method has been shown to be effective in a variety of applications (Teh et al., 2006, 2007; Wang et al., 2013; Blei et al., 2017).

#### 2.5.1 Collapse over hidden variables

To perform a CVB method, we first marginalize over the hidden random variables *μ*_*igk*_’s and *λ*_*gk*_’s. This step also allows us to account for all the variables in a fully Bayesian approach rather than seeking the optimization from the posterior density. By collapsing the joint posterior distribution into a simpler form, we can more efficiently compute the evidence of lower bound and estimate the posterior distribution of the model parameters.

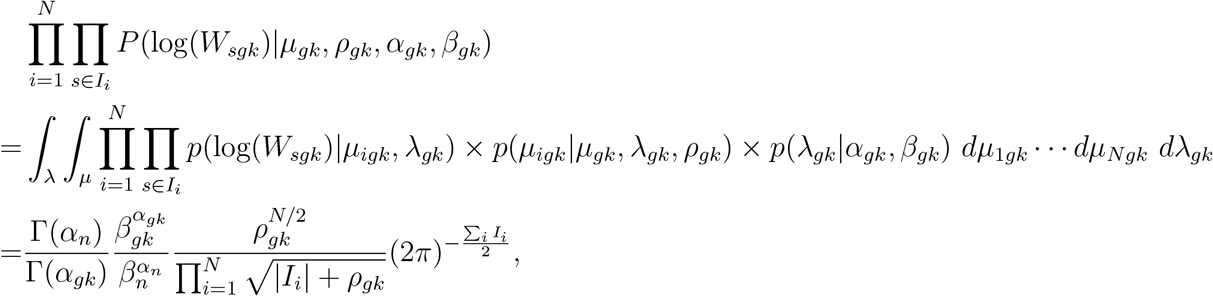

Where 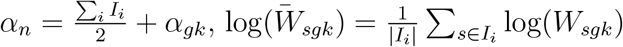, and

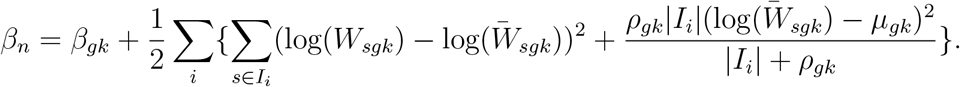

#### 2.5.2 Variational parameters

After we integrate out the latent variables, we introduce the following variational distributions on the remaining unobserved variables, that is *W*_*sgk*_ and *h*_*sk*_,

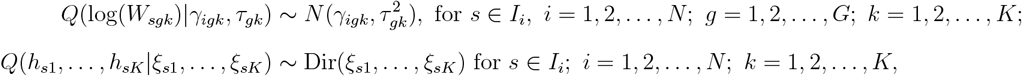

where *Q*(·) denotes the variational distribution that we use to approximate the true posterior density and {*γ*_*igk*_}_*i,g,k*_, {*τ*_*gk*_}_*g,k*_, and {*ξ*_*sk*_}_*s,k*_ are variational parameters that seek to be optimized. In total, there are 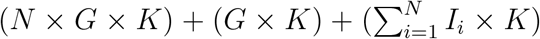 parameters to estimate.

#### 2.5.3 Derivation of the evidence lower bound

Let *Z* = (*W, H*) denote the unobserved variables of interests and ***θ*** = (***α, β, ρ, μ, λ***) denote the hyperparameters. One can approximate the posterior distribution *P* (*Z*|*Y*, ***θ***) by the given variational distributions *Q*(*Z*) by minimizing the Kullback-Leibler (KL) divergence. This is equivalent to maximizing the evidence lower bound (ELBO) in equation 2.3, and the ELBO can be represented by the following five components:

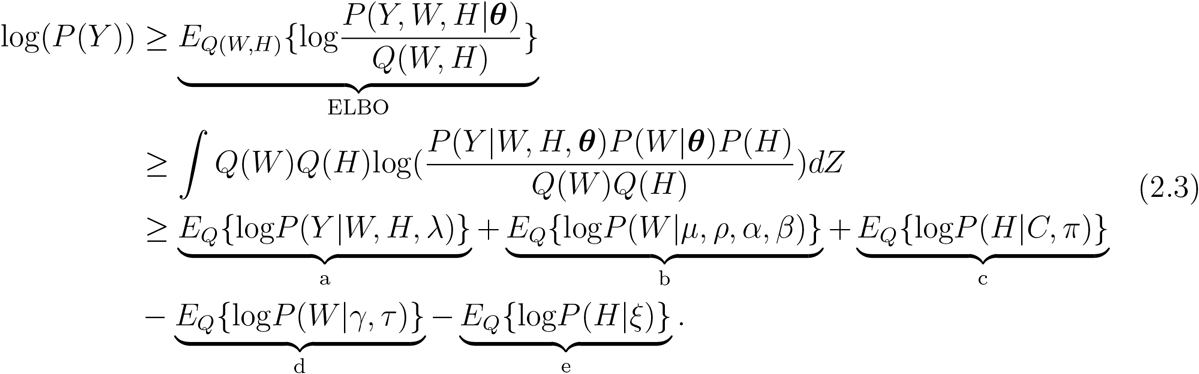

We apply Limited-memory BFGS (Broyden–Fletcher–Goldfarb–Shanno) to iteratively maximize the objective function defined in equation 2.3 and its gradient with respect to variational parameters has been derived to speed the optimization. The stopping criterion is determined if the change in the variational parameters of ELBO between two consecutive iterations is less than some threshold *δ*_*ϵ*_ (default value is 1 × 10^−4^). The analytical calculations of the objective functions and their gradients are presented in the Supplementary Material. An overview of the optimization algorithm of the CVB method is presented in Algorithm 1.

#### 2.5.4 Empirical solution through moments

The relative cell type abundance across all samples can be estimated using the variational parameters obtained through the optimization process. Specifically, for cell type *k* and sample *s* within patient *i*, the estimated proportion of cell type *k* can be calculated as 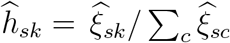, and correspondingly, the overall cell type mean proportions can be obtained as 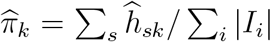. To obtain the intratumor heterogeneity parameter *C*_*i*_ in equation 2.2 for a specific subject *i*, we compute the first and second central moment for *s* ∈ *I*_*i*_ as follows,

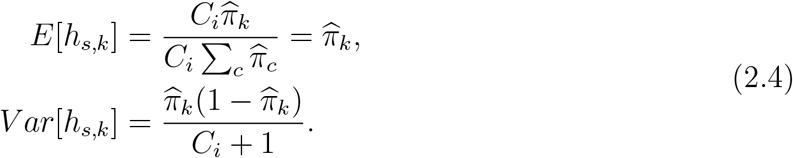

##### Algorithm 1

Collapsed Variational inference algorithm to optimize ICeITH model.

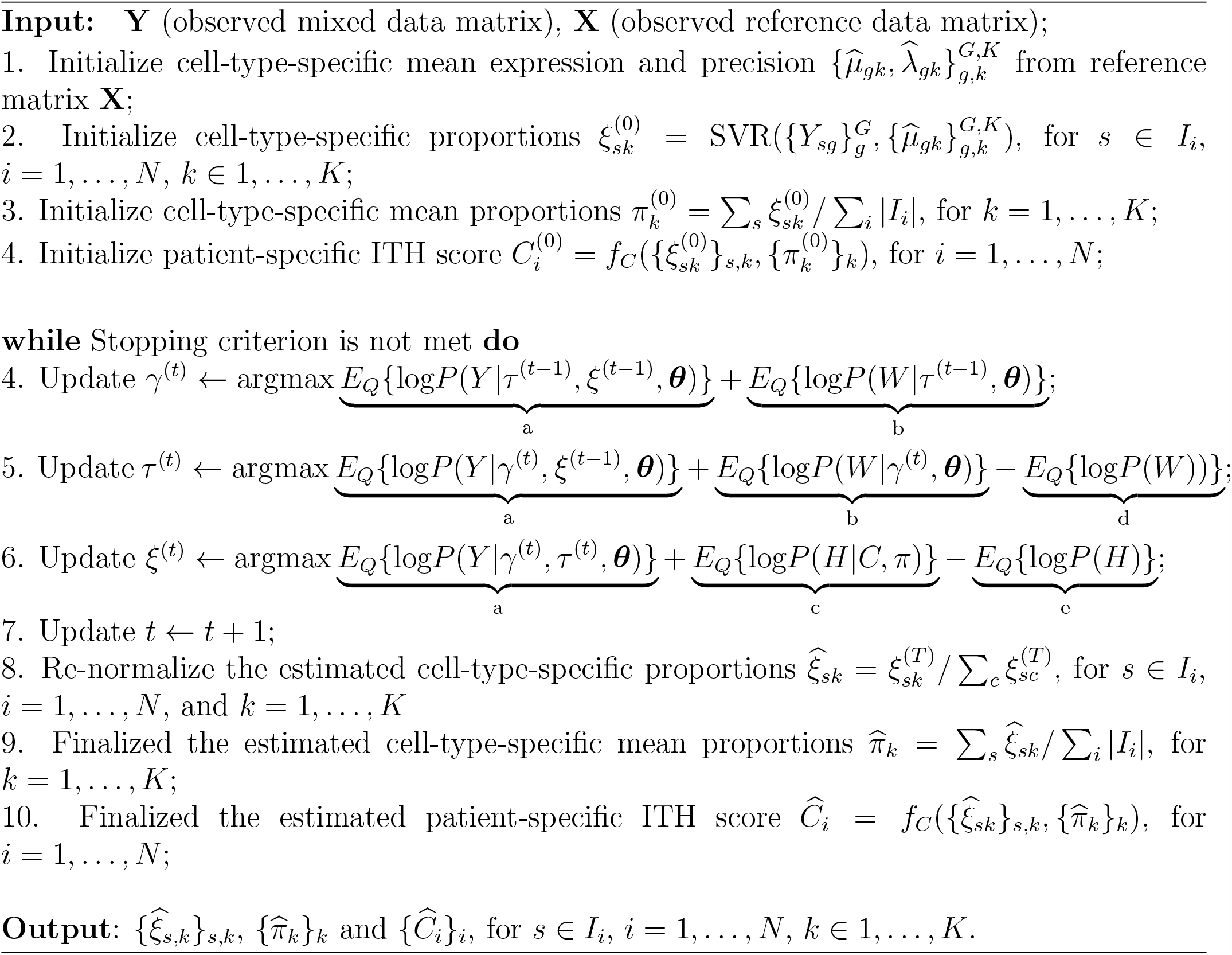

Then, the total variance across samples within subject *i* is 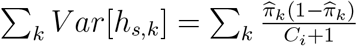, which the ITH score can be calculated as follow:

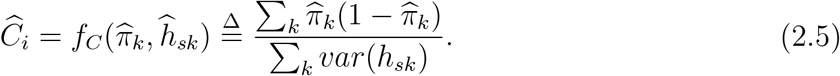

This equation estimates the degree of ITH within each patient based on the variability of the cellular compositions across samples. In particular, the numerator in equation 2.5 represents the expected variance of the cell type proportions across all samples, while the denominator represents the actual variance within each subject. In cases of small sample sizes, it’s possible for the empirically estimated value 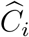 to become negative, even though *C*_*i*_ should always be non-negative theoretically. Nevertheless, the primary interest typically lies in the relative ranking of *C*_*i*_ values, as it provides valuable insights into the association between intratumor heterogeneity and clinical phenotypes.

It should be noted that the estimation of *C*_*i*_ involves summation over all cell types, allowing us to investigate the variability of cellular composition within a targeted subset of cell types, denoted by *T*, where *T* = {*t*_1_, …, *t*_*J*_ } ⊆ {1, …, *K*} is a subset list of all cell types. This allows for the investigation of the degree of ITH within specific subpopulations of cells, which may be of particular interest in certain contexts. Overall, the estimation of cell type proportions and the ITH parameter provides important insights into the composition of the tumor microenvironment and may aid in the development of personalized cancer therapies.

### 2.6 Deconvolution methods for benchmarking

CIBERSORT (Newman et al., 2015) is a deconvolution method that aims to estimate the cell-type-specific composition using *v*-support vector regression, a machine learning method trying to find a hyperplane to dichotomize binary groups by maximizing the margin derived from support vectors (Schölkopf et al., 2000). SVR tolerates data points falsely falling into a constant margin *ϵ* that further decides a *ϵ*-sensitive loss function as below:

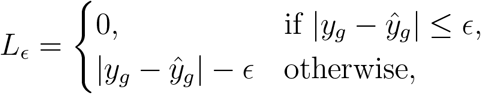

where *y*_*g*_ is the vector of the observed gene expressions for gene *g* and *ŷ*_*g*_ is its expected value. *y*_*g*_ can be decomposed as linear combination of cell-type-specific expressions, which is Σ_*k*_ *w*_*gk*_*h*_*k*_. The loss function can be further defined as:

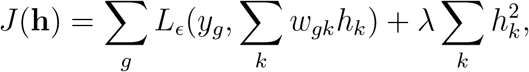

where *λ* is a tuning parameter for the penalty of the regression coefficient, which is cell type abundance in this case. However, choosing a proper *ϵ* tube is difficult in practice. As an alternative solution, CIBERSORT applies *v*-SVR, which asymptotically selects *v* proportions of genes as support vector, with a linear kernel to solve the loss function and select the best results from three tuning values of *v* = {0.25, 0.5, 0.75}. We implement CIBERSORT using the function provided by the website (https://cibersortx.stanford.edu/).

EPIC (Racle et al., 2020) applies a weighted regression-based method to perform deconvolution, the loss function can be defined with the same notation above as follows:

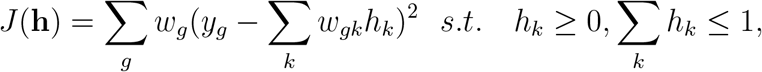

where *w*_*g*_ is the weight for gene *g*. In EPIC, these weights are given by *w*_*g*_ = min(*u*_*g*_, 100 × median(*u*_g_)), where 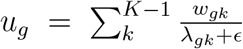 and *λ*_*gk*_ represents the cell-type-specific variance. Therefore, it increases the contribution of signature genes with lower variability to the model fit. Finally, it quantifies unknown cell types by 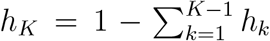. In this paper, we implement the EPIC model by R package EPIC downloaded from GitHub (https://github.com/GfellerLab/EPIC).

## 3 Simulation Study

We conduct extensive simulation studies to evaluate the performance and robustness of our proposed method, ICeITH, as well as benchmark it against CIBERSORT and EPIC. We evaluate the accuracy of estimated cell-type-specific proportions by comparing them to the true proportions using Spearman’s correlation coefficients and root mean squared error (RMSE). Additionally, we assess the ability of each method to discriminate patient stratification by their intratumor heterogeneity score using the F1-score, which combines precision and recall in a harmonic mean (Sasaki et al., 2007)

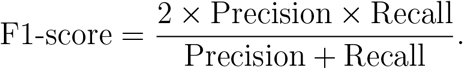

The precision is defined as the ratio of true positives to the total number of patients classified as high intratumor heterogeneity (true positive + false positive). It measures the accuracy of each method in correctly classifying patients into the high intratumor heterogeneity subgroup. The recall is the ratio of true positives to the total number of patients who actually have high intratumor heterogeneity (true positive + false negative). It measures the method’s ability to capture all patients with high intratumor heterogeneity. A higher F1-score indicates better performance in discriminating patients based on their intratumor heterogeneity score. We used this metric to compare and evaluate the effectiveness of each method in patient stratification.

### 3.1 Simulation set-ups

To generate cell-type-specific reference profiles, we first randomly assign 500 genes to four immune cell types. Each gene is assumed to be highly expressed in one of the immune cell types and has lower expression in other cell types. To mimic the heterogeneity seen in real data, we divide the genes into three levels of expression: low, medium, and high. The average expression profiles for each level are simulated from a uniform distribution with distinct ranges. For example, lowly expressed genes have means (under log-scale) ranging from 2 to 4, and the means of cell-specific genes are up-regulated to the range of 3.5 to 5. Similarly, genes with means drawn from the support between 4 to 6 and 6 to 8 represent median and high-level expressed genes, respectively, and the means of cell-specific genes are up-regulated to the ranges of 5.5 to 7 and 7.5 to 9. To capture the mean-variance relationship seen in real data, we calculate the mean-variance relationship from FPKM-UQ normalized data across cell-type-specific genes, assuming genes that share the same average expression profile have the same variability within each cell type. We then generate fifty purified reference samples for each cell type by log-normal distribution with gene-specific average expression profiles and variability, following the strategy from Wilson et al. (2020).

In the second step of the simulation study, we aim to simulate mixed tumor samples for 100 patients with multiple tumor samples assigned to each patient. To better mimic the data characteristics of the real scenarios, we consider including within-subject correlation in multiregion gene expression data generation (Figure 1). Patient-specific average expressions for each cell type and gene are generated by a normal distribution with cell-type-specific mean expression and variance determined at the population level, as per the previous step in the process. We then simulate cell-type-specific expression from a log-normal distribution across samples for each patient. To simulate the remaining immune cell proportions, we first draw the intratumor heterogeneity score from a uniform distribution ranging from 0.1 to 3.0. Next, we simulate cell proportions from a Dirichlet distribution, with the average abundance across patient samples ranging from 0.1 to 0.4. The ultimate expression profile for each sample is obtained as a weighted summation of cell-type-specific expressions.

All the results presented are summarized over 100 Monte Carlo datasets. Overall, this simulation design allowed us to investigate the performance and robustness of our method, ICeITH, under realistic and diverse scenarios of intratumor heterogeneity and cell-type-specific gene expression variability.

### 3.2 Simulation results

#### 3.2.1 Results with four cell types

Based on the simulation settings in the previous section, where the independence assumption between mixed samples within each patient no longer holds, the ICeITH model provides the most accurate estimation of cell type proportions in terms of the Spearman correlation, based on the sampling distribution obtained from 100 Monte Carlo datasets (Figure 3 (a)). The medium value of Spearman correlation equals 0.97, compared with the medium value of 0.92 and 0.88 for CIBERSORT and EPIC, respectively. This indicates that the estimates from ICeITH model not only show a linear relationship with the true values but are also consistent with the rank, reducing the risk of falsely recognizing the dominant cell type. Additionally, the ICeITH model demonstrates its estimation accuracy by investigating the root-mean-square error (RMSE) in Figure 3 (b). Although the sampling distributions of RMSE, obtained from ICeITH and CIBERSORT, seem to be comparable, the medium value (0.045) of RMSE from ICeITH is still better than the medium value (0.049) from CIBERSORT. However, EPIC suffers from the inflation of the estimation variance and false detection or deletion of certain cell types. We further pick an example from simulated datasets to visualize the performance of these three methods (Figure 3 (d-f)). Given the estimated cell type proportions, we empirically estimate ITH scores for these three methods based on equation 2.5 and further dichotomize patients into high ITH and low ITH groups by the medium value of estimated ITH scores. To quantify the classification performance in terms of precision and recall, we further compare the F1-score of all methods (Figure 3 (c)). All three methods provide reasonable F1-score, though our method performs better (medium = 0.78). Additionally, we assess the classification accuracy in terms of Area under the receiver operating characteristics curve (AUC) and ICeITH achieves the best performance (medium = 0.85) (Figure S1 (a)). Cumulatively, we conclude the proposed empirical solution is sufficient to detect patients with different variability of cell type compositions.

**Figure 3.**
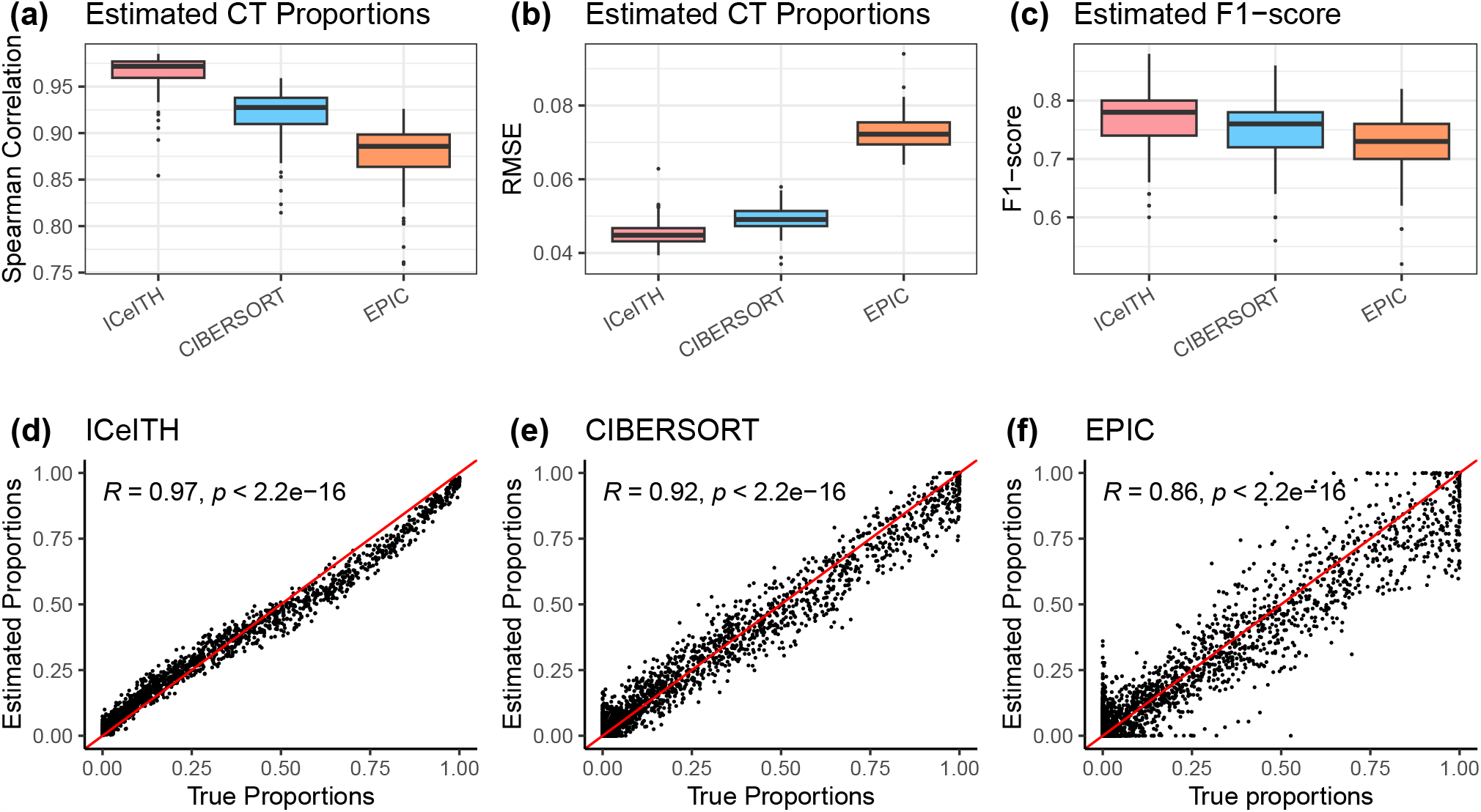
Results of analyzing the simulation data in the setting with four cell types and randomly generated sample numbers. Simulation results of Spearman correlation (a) and Root-mean-square error (RMSE, b) between four true cell type (CT) proportions and estimated CT proportions from three different methods: ICeITH, CIBER-SORT, and EPIC. (c) Sampling distribution of F1-score on low vs. high ITH (intratumor heterogeneity) group, dichotomized by the median value of ITH, from three different methods. Each sampling distribution contains 100 replicates. (d-f) The comparison of four true CT proportions and estimated proportions by three methods with Spearman’s rank correlation test.

#### 3.2.2 Results with 22 cell types

To understand the performance of all methods with substantially more cell types, we also extend our simulation design to 22 cell types with 1000 genes from 50 patients. The rest of the simulation setting follows exactly as discussed in Section 3.1. Figure S2(a) shows that ICeITH provides the highest Spearman correlation (medium 0.63) with the true proportions, while CIBERSORT (medium = 0.37) and EPIC (medium = 0.27) perform poorly due to their failure to identify the correct rank for cell type proportion estimation with the increased number of cells. ICeITH also achieves the lowest RMSE values (medium = 0.04) compared to CIBERSORT (medium = 0.06) and EPIC (medium = 0.08) in Figure S2(b). The inflation of the standard error of the sampling distribution of RMSE in EPIC leads to the false detection and ignorance of certain cell types, as confirmed by examining a specific example (Figure S2(e-g)). We also evaluate the classification performance of all three methods based on the estimated cell type proportions, with ITH scores calculated using equation 2.5. The patients are further classified into high and low ITH groups based on the medium value, and the classification performance is evaluated using the F1-score and AUC. All three methods achieve reasonable F1-score and AUC, with our method performing better (mean F1-score = 0.74, mean AUC = 0.81) (Figure S2 (c,d)). Therefore, our proposed empirical solution can handle large-scale datasets with multiple cell types and is robust to discover the variability of cell type compositions within each patient.

#### 3.2.3 Sensitivity analysis

To evaluate the robustness of the ICeITH model, we conduct a sensitivity analysis by simulating four different scenarios. For each scenario, the same number of samples, from three to six, are assigned to each patient since sequencing above six samples per patient is usually not practical, while other simulation settings remain unchanged. Comparing the sampling distribution of the Spearman correlation from the estimated cell type proportions across three methods (Figure 4 (a)), the ICeITH model outperforms other methods and achieves better accuracy in terms of both mean and standard error with an increased number of samples per patient, while the medium values of the Spearman correlation from replicated experiments remain stable under both CIBERSORT and EPIC. This is because both CIBERSORT and EPIC perform deconvolution independently across samples, while our proposed method takes the inner correlation of samples within each patient into consideration. Although the sampling distribution of RMSE from the ICeITH model suffers from inflation of standard error in the first scenario where patients shared a limited sample size, it rapidly decreased as more samples are available for each patient (Figure 4 (b, d)). For the F1-score derived from the estimates of ITH scores, our method remains comparable with the other two methods when patients suffer from a limited sample size, but it starts outperforming as the sample size for each patient increases (Figure 4 (c)). We observe a similar trend for the AUC across various scenarios (Figures S1 (b)). More importantly, under the least favorable case, where each patient only has limited samples, all three methods still provide a reasonable F1-score. This indicates that our method is robust for calculating the relative cell type abundance and detecting patients with different variability of cell type compositions

**Figure 4.**
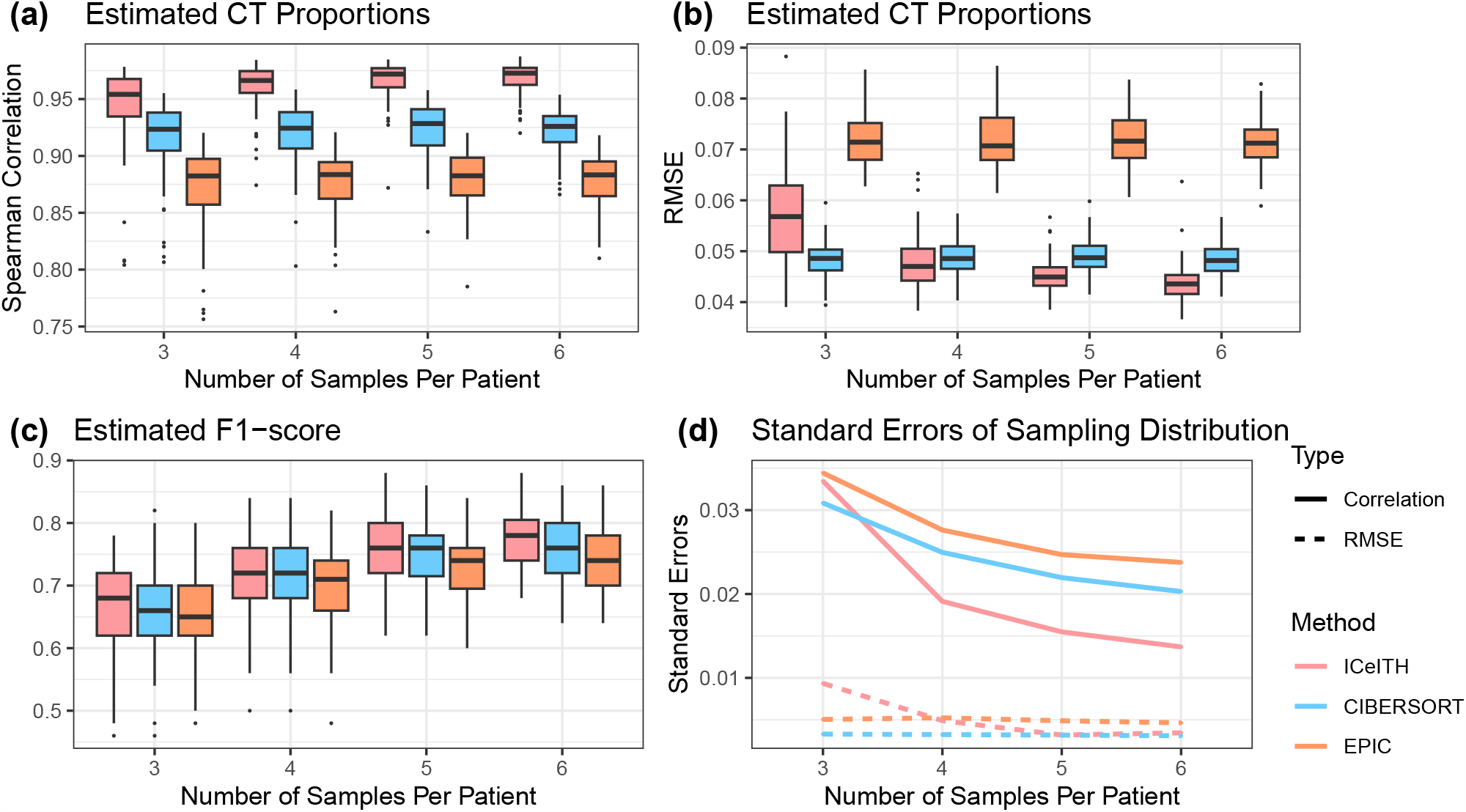
Results of analyzing the simulation data in the setting with four cell types and fixed sample numbers. Sampling distribution of Spearman Correlation (a) and RMSE (b) between true and estimated CT proportions from three different methods: ICeITH, CIBERSORT, and EPIC based on the scenarios where every patient shares the same number of samples. (c) Sampling distribution of F1-score generated from the estimated ITH scores by each method on each scenario. (d) Standard errors of sampling distribution from (a) and (b) across different methods and scenarios.

In addition, we evaluate the robustness of our proposed model in scenarios (1) where genes from the reference panel exhibit different expression patterns compared to the observed samples and (2) under varying degrees of within-subject correlations. ICeITH maintains a superior performance compared to CIBERSORT under these conditions (Figure S3,4). Details are provided in the Supplementary Material Section S3.3.

## 4 Real data application

### 4.1 Reference profile

To construct the reference profile, we download a gene expression dataset from the NCBI Gene Expression Omnibus with accession number (GSE60424, Linsley et al. (2014)). This study investigates the whole transcriptome signatures of six immune cells and whole blood from patients. While the whole blood is collected into a Tempus tube, the rest of the primary fresh blood sample is processed into highly purified populations of neutrophils, monocytes, B cells, CD4 T cells, CD8 T cells, and natural killer cells. RNA extraction is performed on each cell subset, and progressed into RNA sequencing libraries (Illumina TruSeq), with a targeted read depth of around 20M reads. The reads were then demultiplexed and mapped to human genome hg19. The reads were summarized into HTSeq counts by the RSEM package and annotated using the Ensembl database GRCh37 (Hubbard et al., 2002).

### 4.2 Application to TRACERx data

#### 4.2.1 Preprocessing TRACERx data

Multi-region WES and RNA sequencing data of the TRACERx cohort are obtained from Jamal-Hanjani et al. (2017). Whole-exome sequencing is performed on extracted DNA samples and described in Jamal-Hanjani et al. (2017). Tumor purity for each sample is estimated by Sequenza (Favero et al., 2015). Some matched samples also proceed to RNA sequencing. Specifically, alignment and reads mapping (human hg19 reference genome) is performed by the STAR package (Dobin et al., 2013) and tabulated to HTSeq counts by the RSEM package (Li et al., 2011). Clinical information for the TRACERx cohort is downloaded from https://www.nejm.org/doi/full/10.1056/NEJMoa1616288.

The mixed tumor counts are then normalized together with reference counts by the TMM procedure using R package edgeR (Robinson et al., 2010), and the batch effect is adjusted using R package combat-seq (Zhang et al., 2020). To ensure comparability across samples, we perform scale normalization with total UMI across samples. We apply signature genes, obtained from RNA-seq reference (LM6) available on the CIBERSORT website (https://cibersort.stanford.edu/download.php), to perform deconvolution. In addition, we filter out signature genes with cell-type-specific variance greater than 1.5, and this results in 523 genes after matching genes with tumor samples. Note: the reference matrix is sparse due to the blood sample type. We add a small number (e.g. 0.01) to the cell-type-specific variance that equals 0 so that it guarantees a feasible optimization when evaluating the gradient of variational parameter *τ* in equation S.9.

#### 4.2.2 Results on TRACERx data

We apply the ICeITH model and CIBERSORT to analyze RNA-seq data in patients with non-small cell lung cancer collected from the TRACERx cohort (Jamal-Hanjani et al., 2017). The multi-region DNA-seq and RNA-seq data are available in 45 patients (patients with a number of regions available greater than one), and it results in 140 tumors in total (Table S1). We seek to associate the survival outcome with intratumor heterogeneity scores estimated from our proposed method.

Specifically, we fit both the ICeITH model and CIBERSORT on the normalized dataset to estimate the relative abundance for six immune cell types. Given the tumor cell fractions obtained by Sequenza (Favero et al., 2015) whole-exome sequencing data for all samples, we obtain the adjusted immune cell type proportions by renormalizing them to one minus tumor cell fraction for each sample.

The estimated proportions from six immune cell types and tumor cells are displayed in Figure 5 (a). We observe a low concentration of most immune cell types, with the exception of a high variation in monocyte infiltration. This result is consistent with CIBSERSORT estimates (Figure 5 (b), Spearman correlation = 0.82). Previous studies (Jamal-Hanjani et al., 2017; Zhang et al., 2014) shed light on the spatial divergence in genomic instability and provide the impact of this intratumor heterogeneity on the increased risk of patient survival. The tumor microenvironment is also dominated by tumor-surrounded interactions (Whiteside et al., 1998), i.e., various immune effector cells with pro- and anti-tumor functions. Therefore, we are interested in investigating the influence of intratumor heterogeneity, defined as the spatial variability of immune cells that provide either pro- or anti-tumor activities within a patient, on disease-free survival probability. For immune cells (e.g., neutrophils and monocytes) act pro-tumor functions, we estimate the pro-tumor ITH scores using adjusted Neutrophils and Monocytes proportions by equation 2.5, and then we classify patients as high and low pro-tumor ITH groups, respectively, based on the medium value. The results show that patients with a higher pro-tumor ITH are at significantly increased risk compared with a lower pro-tumor ITH (Figure 5 (c), *P* value = 0.05), which indicates a greater spatial divergence of immune cells functioning as a pro-tumor process will suppress the immune response to tumors and negatively affect patient survival. However, it appears that this finding is confounded by the overall immune context of each patient. This is motivated by the findings of AbdulJabbar et al. (2020) that differentiate highly immune-infiltrated tumor regions by the number of immune hot and cold tumors at a population level reveal patient survival. We further stratify a subset of patients (*n* = 44) with available immune identification into immune context low vs. high groups based on the median number of immune cold and hot regions. Interestingly, the most at-risk group is patients with a freeze immune context (i.e., low immune context and low pro-tumor ITH), and the other groups stayed at a comparably decreased risk (Figure 5 (e), *P* value = 0.003). Meanwhile, tumors with increased ITH obtained from immune cells (i.e., B cells, CD4T, CD8T, and natural killer cells) shaped as anti-tumor activities seem to be associated with decreased risk (Figure 5 (d), *P* value = 0.12), which indicates immune cells with anti-tumor process play an important role on eliminating tumor cells. This is further proven when we stratify patients by both immune context and anti-tumor ITH. We observe patient with a increased anti-tumor ITH is always associated with a decreased risk of survival within each immune context (Figure 5, *P* value = 0.003). Cumulatively, we conclude that patients stratified by both immune context and ITH further provide us further evidence that the variability of immune cell type compositions with different activity processes is capable of predicting the prognosis in early-stage lung cancer (Figure 5 (e,f), *P* value = 0.003).

**Figure 5.**
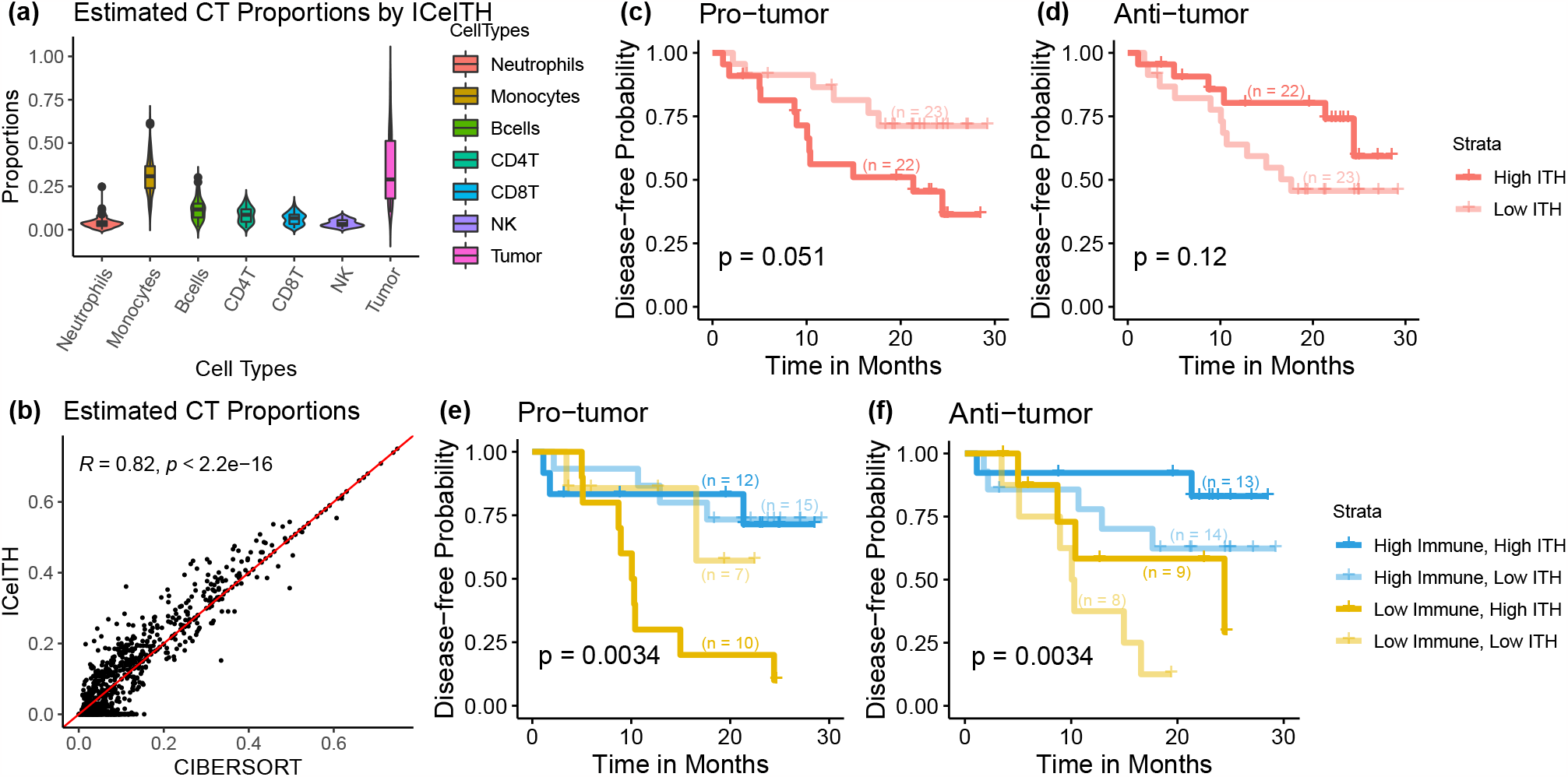
Results of analyzing the TRACERx data. (a) Distribution of relative abundance for six immune cell types and tumor cells estimated by ICeITH and Sequenza, respectively. (b) Dot plot to compare the estimates between the ICeITH model and CIBERSORT. Kaplan-Meier (KM) curves for disease-free probability with high vs. low ITH estimated under (c) pro-tumor and (d) anti-tumor process immune cells for 45 patients from the TRAC-ERx study. KM plots for disease-free probability with high vs. low ITH estimated under pro-tumor and (f) anti-tumor process immune cell, stratified by different immune levels, for 44 patients in the TRACERx study. The sample size for each stratification is marked on the KM curve.

Motivated by the simulation results that the performance of the ICeITH model increases as more samples are sequenced from each patient, we further keep patients (*n* = 28) with more than two regions available and explore the association of intratumor heterogeneity of targeted immune cell types to disease-free survival outcomes. We observe a similar trend as shown in Figure 5. Specifically, for immune cells (i.e., neutrophils and monocytes) that have pro-tumor functions, patients with a higher ITH are at a significantly increased risk compared with a lower ITH (Figure S5 (a), *P* value = 0.03). Data from patients stratified by both immune context and ITH further confirms that a higher ITH are associated with a worse outcome (Figure S5 (c)). However, possibly due to the limited sample size, such an effect is not statistically significant (*P* value = 0.1). Meanwhile, patients with high ITH in anti-tumor immune cells (i.e., B cells, CD4T, CD8T and natural killer cells) are at a reduced risk of survival outcome (Figure S5 (b), *P* value = 0.04). When we further stratify patients with overall immune context, patients with high ITH still consistently have better survival outcomes (Figure S5 (d)). Subsetting patients with a larger sample size provides additional evidence that our proposed method is able to capture the targeted immune cell variability and thus may predict prognosis in early-stage lung cancer.

Additionally, we use summary statistics of immune cell prevalence at the patient level, such as quartiles, median, mean, and maximum, to categorize patients into different risk levels as an alternative approach to our proposed ITH score. The results indicate that while neutrophil cell prevalence did not strongly impact survival, increased heterogeneity in monocyte cells was associated with poorer survival outcomes (Figure S6).

### 4.3 Application to MDA-ITH data

We further extend our analysis to investigate the impact of ITH on the prognosis of lung cancer patients from the MD Anderson-ITH (MDA-ITH) project, where multi-region RNA sequencing technology is performed on 25 patients, and it results in 116 samples in total (Table S2). We normalize the mixed bulk counts together with reference counts and follow the same steps as described in the Section 4.2.1. After we filter out signature genes with cell-type-specific variance greater than 1.5 and match genes with mixed bulk samples, the data results in 605 genes to perform deconvolution. Previous MDA-ITH studies mostly focused on the ITH of mutations and copy number variations, but the heterogeneity of immune cell infiltration has not been carefully studied before. Again, we apply the GSE60424 dataset (Linsley et al., 2014) as a reference profile and fit both the ICeITH model and CIBERSORT on the normalized dataset followed by the same step from the previous section. Due to the lack of matched DNA-seq data, we are not able to adjust estimated immune cell type proportions by tumor fraction for each sample. And thus, mixed samples sequenced from each patient consist of both tumors and healthy tissue. This may give us less power to detect the association between ITH and overall survival outcomes.

The estimated proportions from six immune cell types are presented in Figure 6 (a). Despite the great variation in the prevalence of neutrophils and monocytes cells, we observe the majority of immune cell types (i.e.g, B cells, CD4T, CD8T, NK) with relatively low abundance across samples. The correlation between the ICeITH and CIBERSORT estimates is lower than TRACRx but still reasonable (Figure 6 (b), Spearman correlation = 0.58). This may partially be due to the fact that mixed bulk samples are sequenced from either tumor or healthy tissues so it adds an extra confounding factor for both deconvolution methods. To further investigate the impact of intratumor heterogeneity shaped by immune cells with different underlying activity processes on overall survival, we first estimate the pro-tumor ITH score from neutrophils and monocyte abundance for all MDA-ITH patients. Patient stratification by pro-tumor ITH level did not show a significant association with survival outcome (Figure 6 (c), *P* value = 0.42). The true signal may be eliminated by the limited number of patients enrolled in the study or a mix of tumoral and healthy tissues sequenced for deconvolution. On the other hand, we observe patients with increased anti-tumor ITH, obtained by immune cells functioning as an anti-tumor process, remain associated with a lower risk of overall survival (Figure 6 (d), *P* value = 0.012). This finding is consistent with results that appear in TRACERx data, and together we draw the conclusion that the variability of immune cell-type compositions can serve to predict the prognosis in early-stage lung cancer.

**Figure 6.**
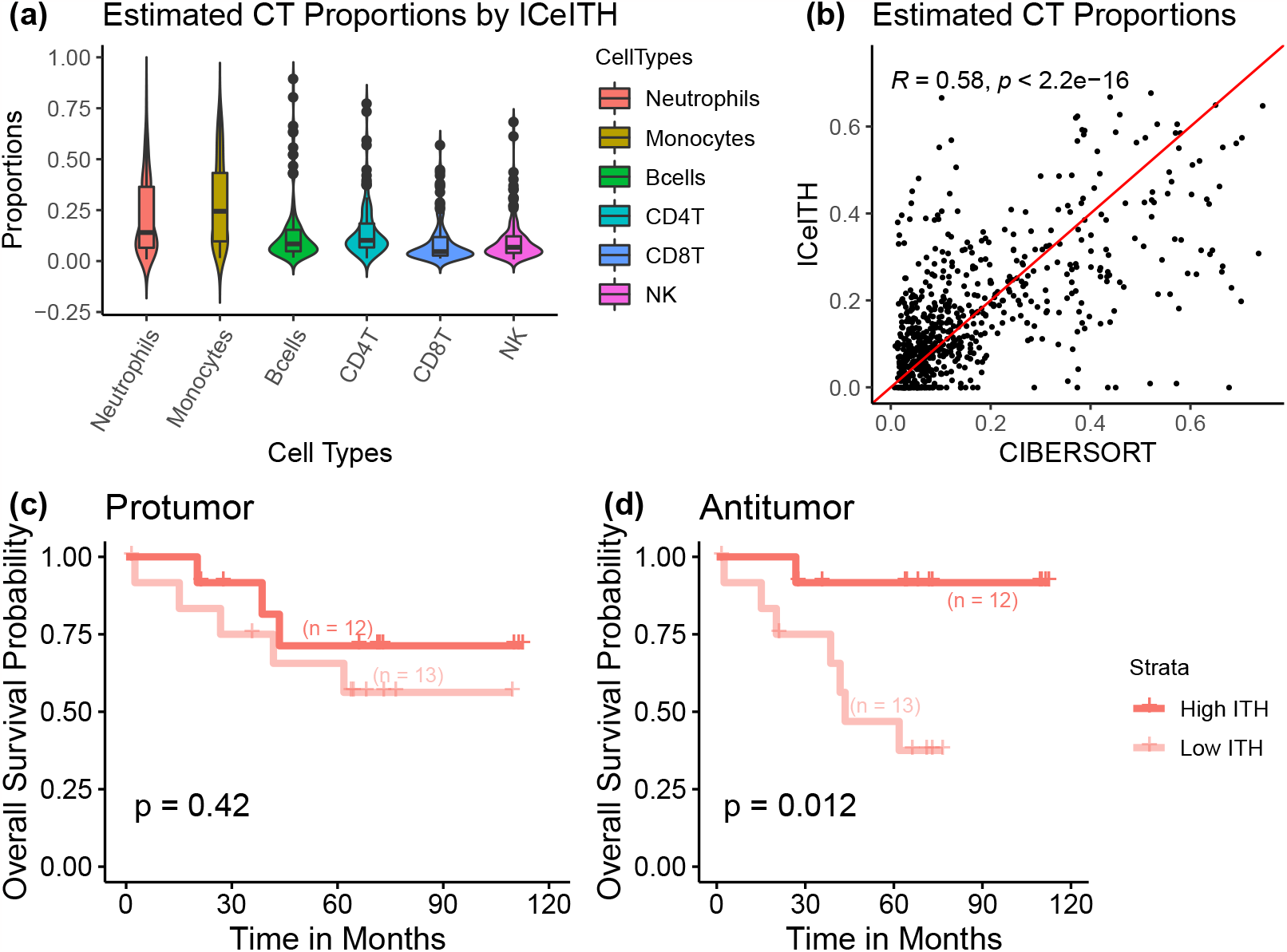
Results of analyzing the MDA-ITH data. (a) Distribution of estimated proportions for six immune cell types by the ICeITH model. (b) Dot plot to compare the estimates between ICeITH model and CIBERSORT. KM curves for disease-free probability with high vs. low ITH estimated by immune cells under (c) pro-tumor and (d) anti-tumor process for 25 patients from the MDA-ITH study. The sample size for each stratification is marked on the KM curve.

## 5 Discussion

In this paper, we propose a novel statistical method, ICeITH, to estimate the immune cell-type abundance from tumor samples using multi-region transcriptomic sequencing data. ICeITH is built on a hierarchical Bayesian approach and has several novelties to current state-of-the-art methods. First, ICeITH models pure cell-type-specific expression profiles as a log-normal distribution to account for the over-dispersion in data with patient-specific mean expression parameters. Our method allows the modeling of patient-specific mean expression across tumor samples from the same patient to capture the within-subject correlations. Second, our proposal adopts a Bayesian approach that is able to leverage the prior information from the reference profile. Compared with previous works that fix the cell-type-specific profiles as a given constant, a Bayesian approach has more flexibility when the cell-type-specific profiles have variations among patients. Lastly, ICeITH assumes the TIL compositions follow Dirichlet distributions with a patient-specific parameter. Such modeling can naturally quantify the variability of targeted cellular composition (e.g. immune cells shape pro- and anti-tumor process) across samples from a single patient for intratumor heterogeneity estimations. We demonstrate that patient stratification based on ITH score is closely associated with patient survival outcomes.

The current real data analysis is limited to the six immune cell types. This is because as the number of cell types increases, the difficulty of accurately estimating the proportions while accounting for the variation within each subject also greatly increases. This is especially true when the number of samples collected from each subject is limited in real-world scenarios. As technology advances and collecting bulk RNA samples becomes cheaper, it will be more feasible to collect larger numbers of samples from each patient. In such cases, we believe our ICeITH model is still capable of handling a higher number of cell types when the sample size is large enough.

The proposed method uses a reference panel to provide prior knowledge on the pure tissue profile and guide the deconvolution process. However, it should be acknowledged that the reference panel and the data of interest are often obtained from different studies. Integrating data across studies can involve complicated platform/sample/technology heterogeneity, introducing substantial analytical bias into the results. Such issues have also been recognized and investigated in recent publications, e.g., Chen et al. (2022) proposed methodologies for integrating multiple reference panels for deconvolution, while Dong et al. (2021) developed an ensemble method that aggregates deconvolution results estimated with different reference panels. These methods specifically tackled the challenge of mitigating cross-study heterogeneity when integrating data from diverse sources.

Our proposed method solves the model efficiently by applying the collapsed variational inference method in the optimization step. We update the variational parameters including patient-specific mean expression and variability and cell type proportions at each iteration. To speed up the optimization, we initialize cell type proportions with SVR and ICeITH converges well under both simulation and real data analyses. We derive closed-form gradients from the objective function to further shorten the computational time. The total running time of the ICeITH model in a dataset containing 140 samples and 523 genes is usually within 30 minutes (using R on a MacBook Pro, 2.4 GHz Intel Core i5).

Currently, our method needs additional information for tumor purity adjustment as our reference panel only consists of immune cell types. Although it is possible for researchers to obtain such information based on existing tools, similar to our analytical procedure with the TRACERx data, it actually provides great convenience if such a step can be automatically incorporated into the analysis pipeline. We acknowledge this limitation and suggest that a comprehensive evaluation of existing tools is beyond the scope of this work. We consider this an important extension for future works. On another thread, our current method applies to multi-region RNA-seq data as motivated by the TRACERx and MDA-ITH studies. In reality, a more common scenario is the multi-omics sequencing of multiple tumor slices for the same patients. The same principle of the proposed method should apply to the multiomics application, but the data distributions will be different. This is another promising direction for future research, and could potentially have a greater impact on ITH research.

## Acknowledgments

This study makes use of data generated by the TRAcking Non-small Cell Lung Cancer Evolution Through Therapy (Rx) (TRACERx) Consortium and provided by the UCL Cancer Institute and The Francis Crick Institute. The TRACERx study is sponsored by University College London, funded by Cancer Research UK and coordinated through the Cancer Research UK and UCL Cancer Trials Centre (Jamal-Hanjani et al., 2017).

This study is supported by the MD Anderson Lung Cancer Moon Shot Program, the Cancer Prevention and Research Institute of Texas Multi-Investigator Research Award grant (RP160668), the National Cancer Institute of the National Institute of Health Research Project Grant (R01CA234629-01), and the UT Lung Specialized Programs of Research Excellence Grant (P50CA70907).

## Supplementary

The Supplementary Material contains additional model derivation and simulation results.

## S1 Model derivation

### S1.1 Derivation of objective function

Five components appear in the evidence lower bound in equation 2 and their derivation with respect to variational parameters can be analytically derived as follows:

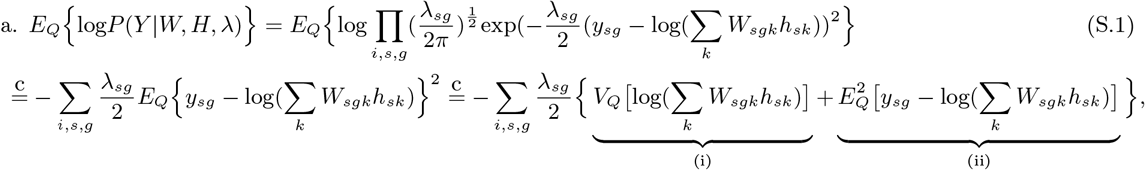

so that

i. 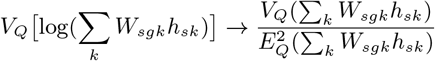, (by delta method), and
ii. 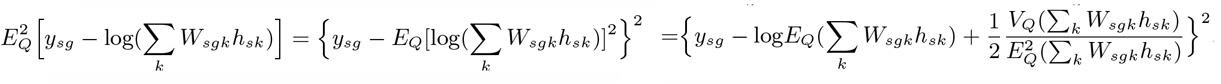, (by second order Taylor expansion), where

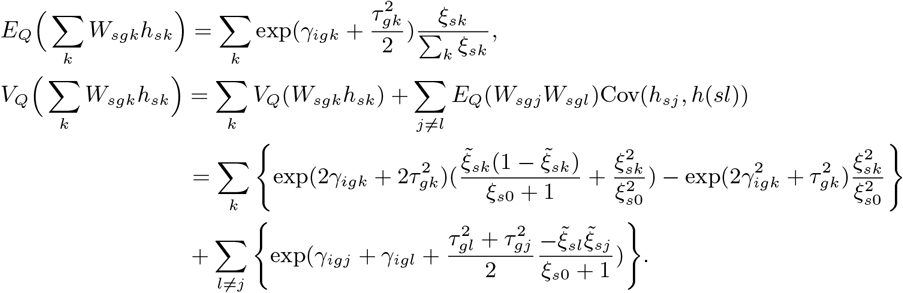

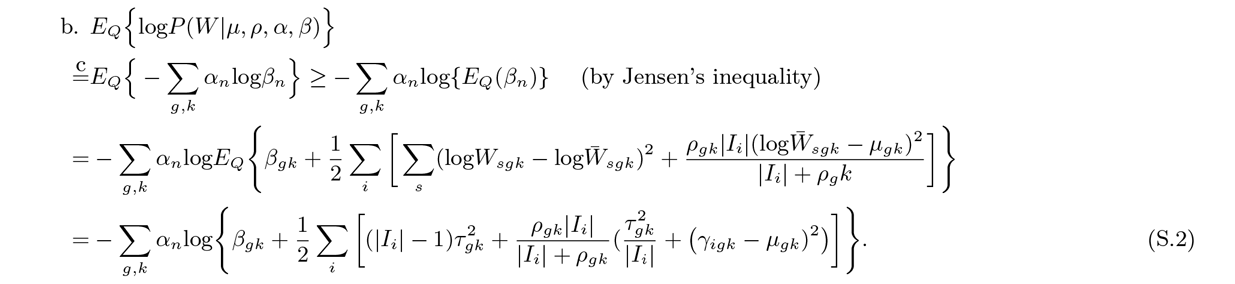

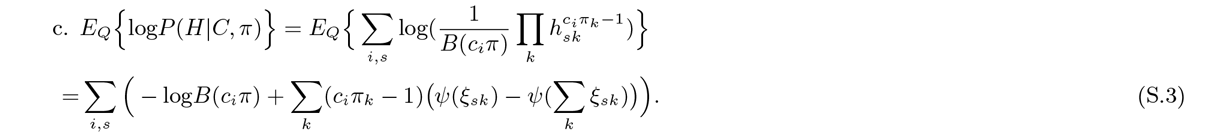

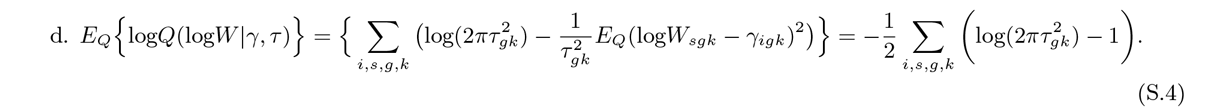

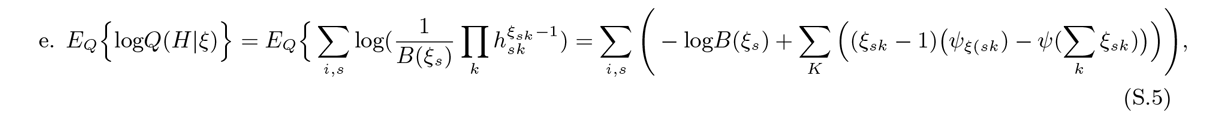

where 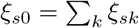 and 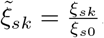.

### S1.2 Derivation of gradient function

#### S1.2.1 Gradient of γ_igk_

The gradient of *γ*_*igk*_ is equivalent to 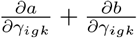, and each term is to be derived as follows:

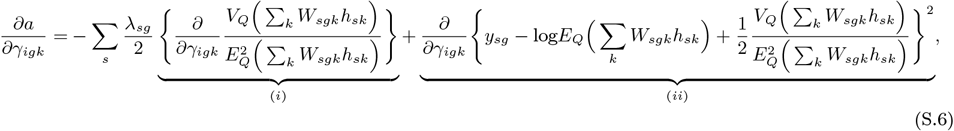

i. 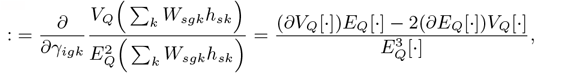
ii. 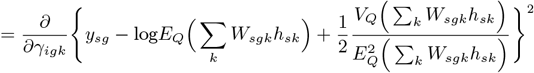

where

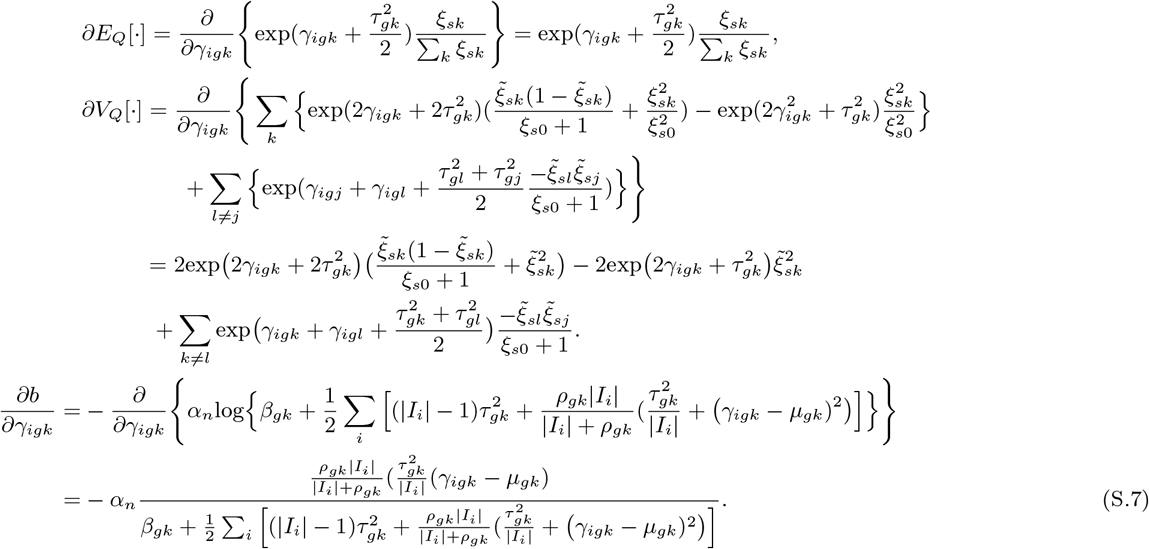

The other terms do not involve *γ*_*igk*_, therefore,

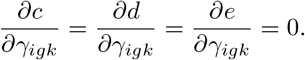

#### S1.2.2 Gradient of τ_gk_

The gradient of *τ*_*gk*_ is equivalent to 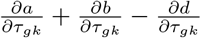, and each term is to be derived as follows:

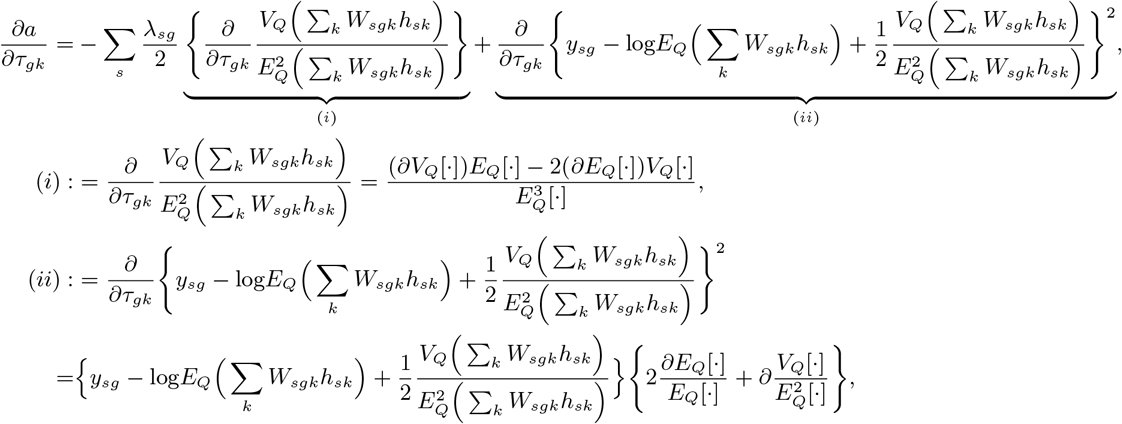

where

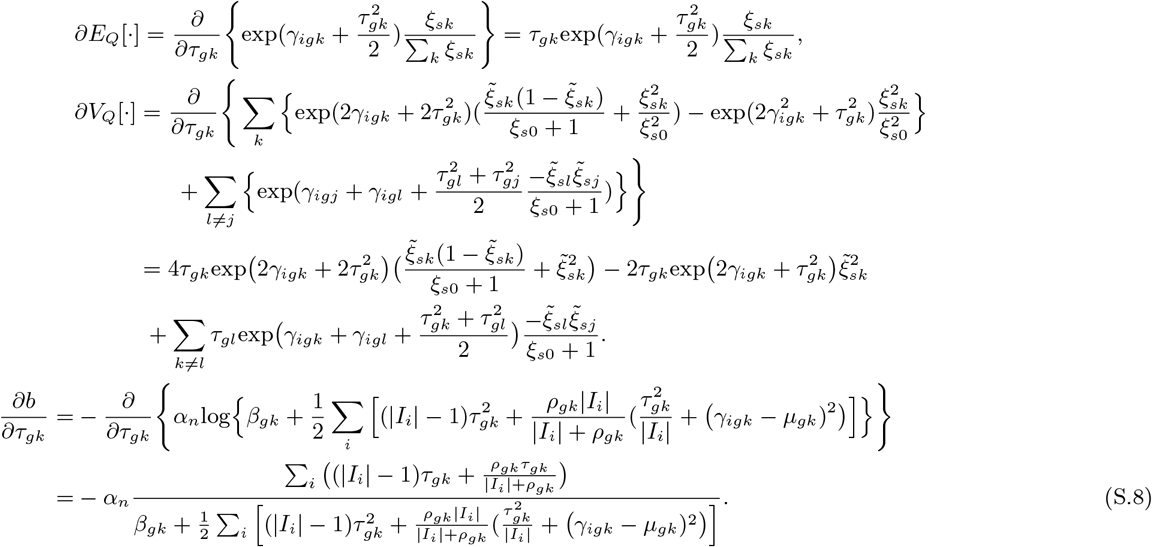

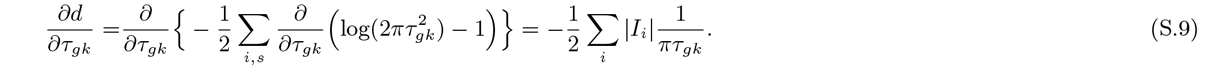

Other terms do not involve *τ*_*gk*_, therefore,

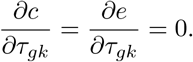

#### S1.2.3 Gradient of ξ_sk_

The gradient of *ξ*_*gk*_ is equivalent to 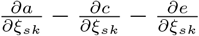, and each term is to be derived as follows (Andrade Barbosa et al. 2021):

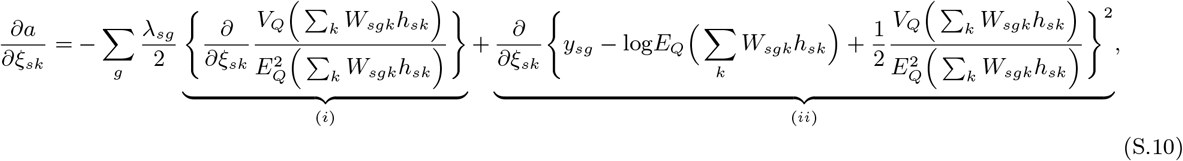

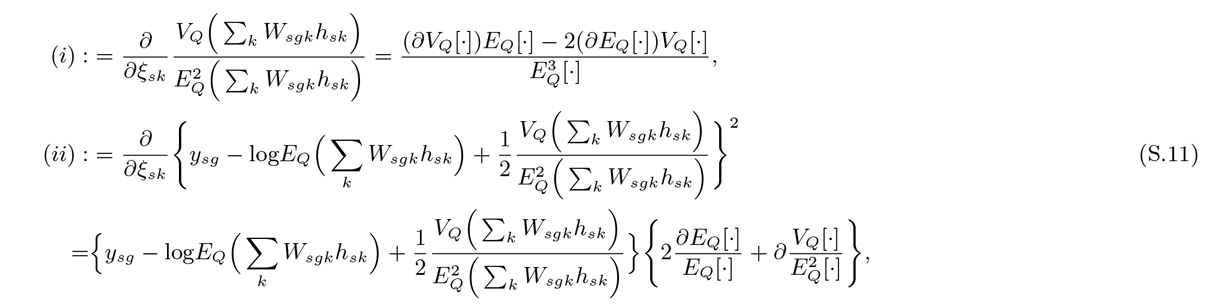

where

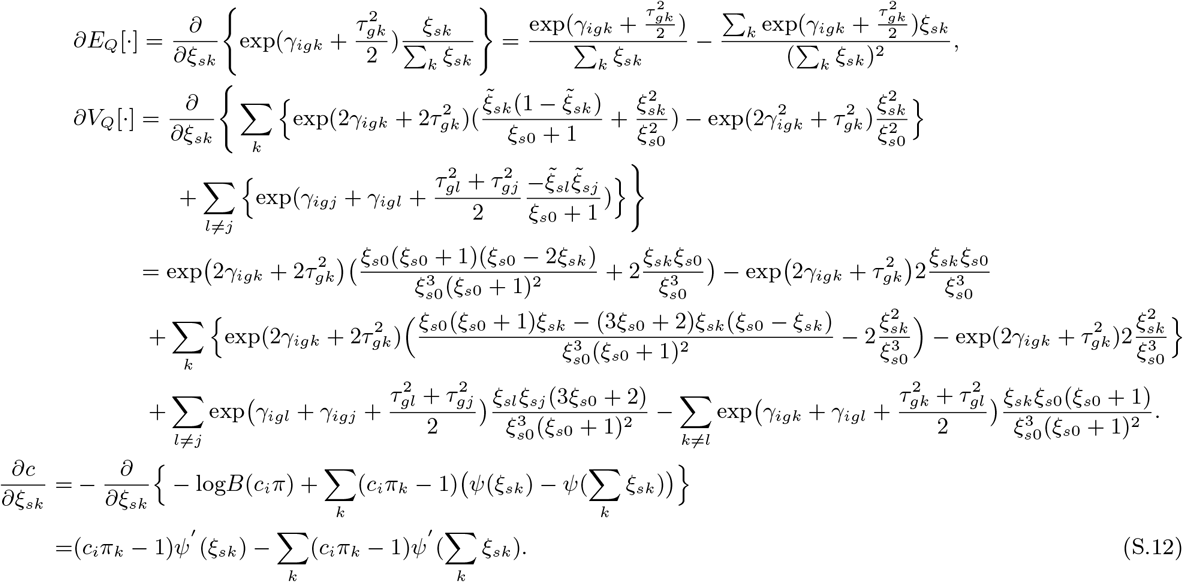

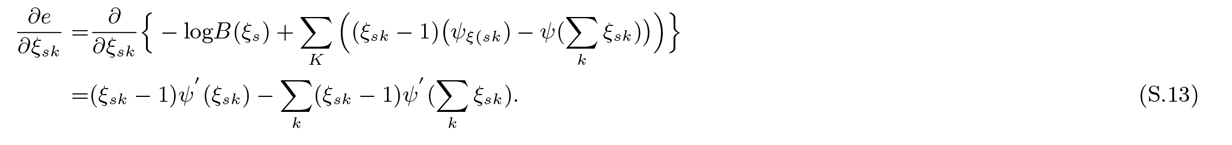

Other terms do not involve *ξ*_*sk*_, therefore,

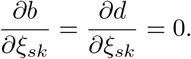

## S2 Overview and implementation of existing methods

### S2.1 Deep learning method to detect lymphocyte

In AbdulJabbar et al. 2020, the author build a convolutional neural network that trained on 5,951 single-cell annotations by pathologists. This approach allows for spatially detect cancer cells, lymphocytes, and stromal cells (fibroblasts and endothelial cells) in H&E-stained images from TRACERx 100 cohort (Jamal-Hanjani et al. 2017). It mainly focuses on differentiating highly from poorly immune-infiltrated tumors by classifying immune hot and immune cold regions based on the percentage of lymphocytes. An immune hot region contains a lymphocyte percentage greater than a quarter s.d. above the median lymphocyte percentage, and an immune cold region contains a lymphocyte percentage below a quarter s.d. of the median lymphocyte percentage. It has been confirmed that patient with a number of immune cold region greater than medium is significantly associated with an increased risk of disease-free survival outcome. We download this clinical information (https://www.nature.com/articles/s41591-020-0900-x) comparable across TRACERx patients to assist our analysis.

## S3 Additional simulation results

### S3.1 Simulating four cell types

**Figure S1:**
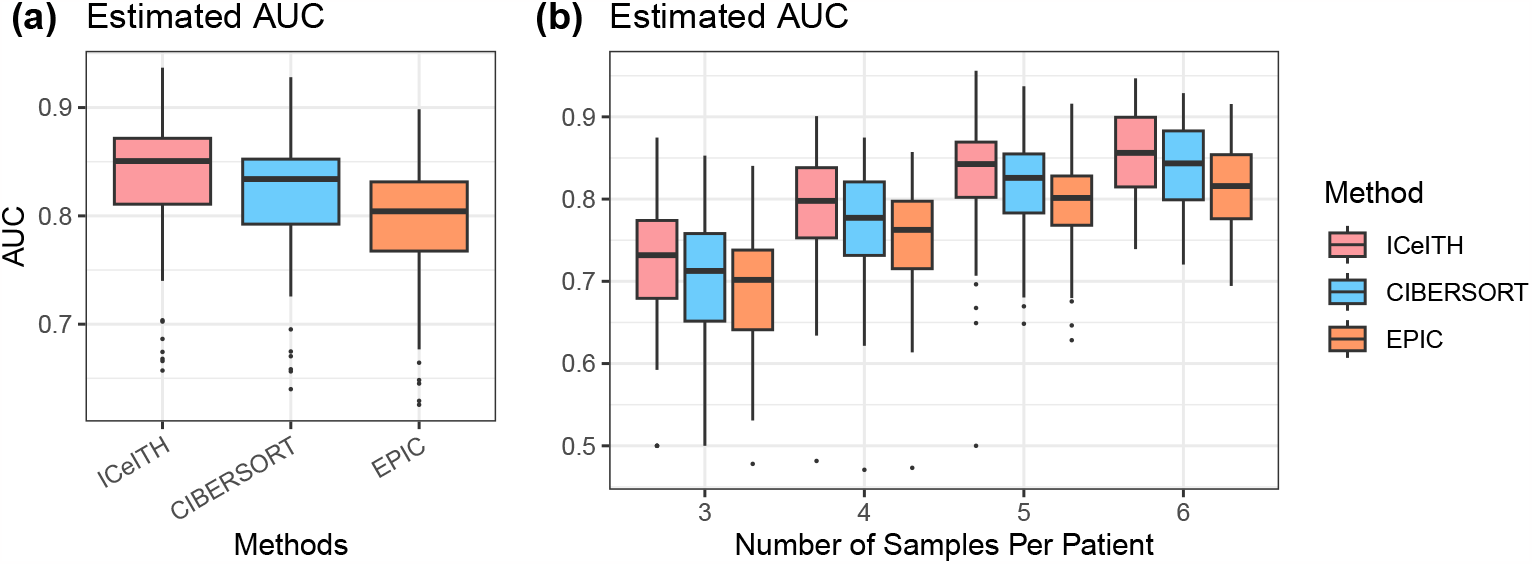
(a) Sampling distribution of AUC (Area under the receiver operating characteristics curve) on low vs. high ITH (intratumor heterogeneity) group, dichotomized by the median value of ITH, from three different methods. (b) Sampling distribution of F1-score generated from the estimated ITH scores by each method on each scenario. Each sampling distribution contains 100 replicates.

### S3.2 Simulating 22 cell types

**Figure S2:**
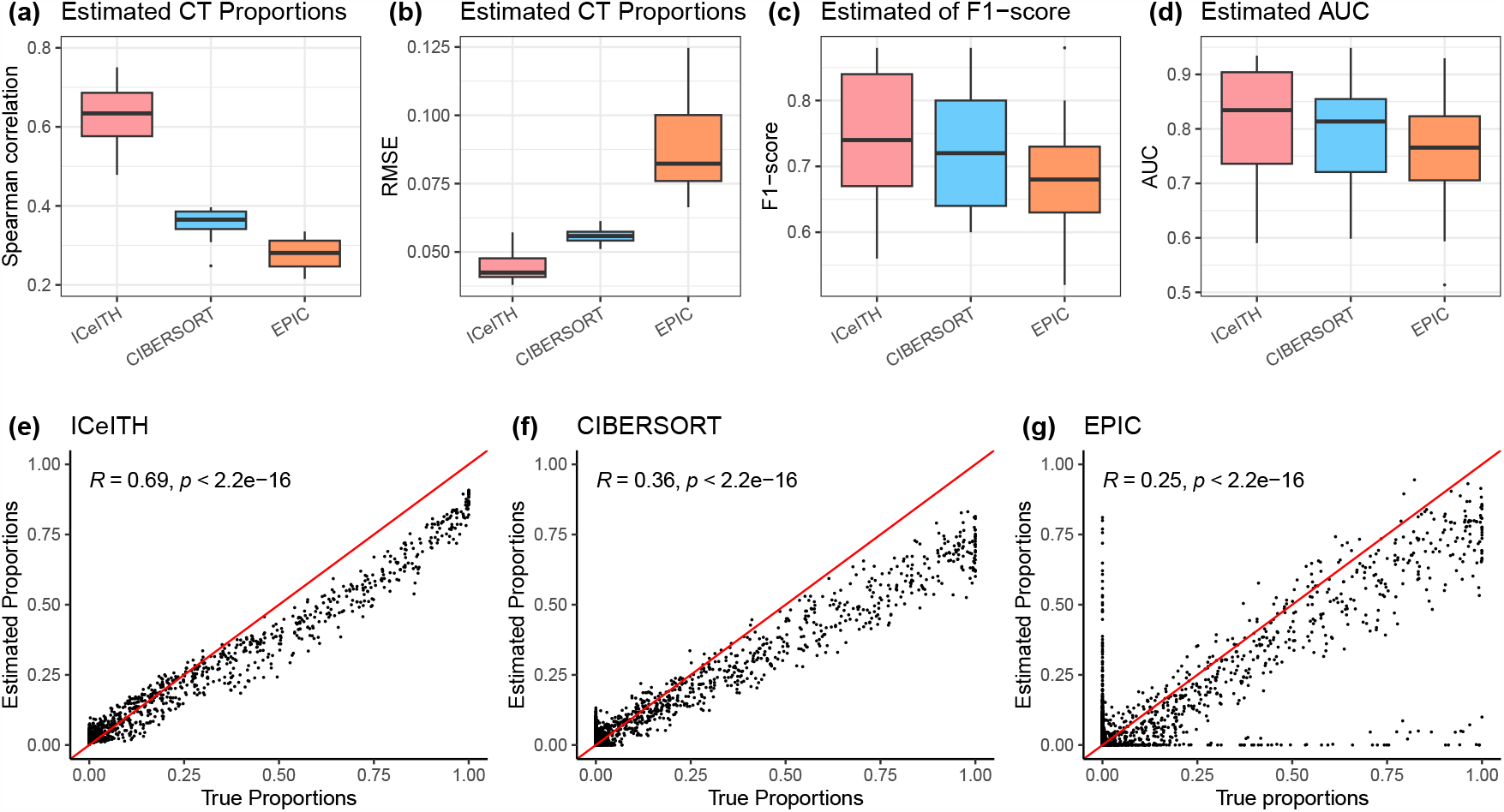
Sampling distribution of Spearman correlation (a) and RMSE (Root-mean-square deviation, b) between 22 true CTs (cell types) proportions and estimated CT from three different methods: ICeITH, CIBERSORT and EPIC. Sampling distribution of F1-score (c) and AUC (d) on low vs. high ITH (intratumor heterogeneity) group, dichotomized by the median value of ITH, from three different methods. Each sampling distribution contains 20 replicates due to computational cost. (e-g) The comparison of 22 true CT proportions and estimated proportions by three methods with Spearman’s rank correlation test.

### S3.3 Sensitivity analysis

We conducted a sensitivity analysis to evaluate the robustness of our proposed model in scenarios where genes from the reference panel exhibit different expression patterns compared to the observed samples. Following the simulation parameters outlined in Section 3.1 in the main text, we introduced a proportion of aberrant genes into the reference profile by adding random Gaussian noise with a mean of 0 and varying levels of variance (0.5 or 1).

Figure S3 presents the Spearman correlations between the true and estimated cell type proportions for both ICeITH and CIBERSORT, across different proportions of aberrant genes and varying levels of Gaussian noise variance. Notably, when the Gaussian noise has low variance (e.g., 0.5), ICeITH consistently outperforms CIBERSORT, maintaining comparable performance across varying proportions of aberrant genes. However, as the discordance within the reference profile increases (e.g., with a higher Gaussian noise variance of 1), both methods exhibit reduced performance as the proportion of aberrant genes grows. Despite this, ICeITH maintains a superior performance compared to CIBERSORT under these conditions.

**Figure S3:**
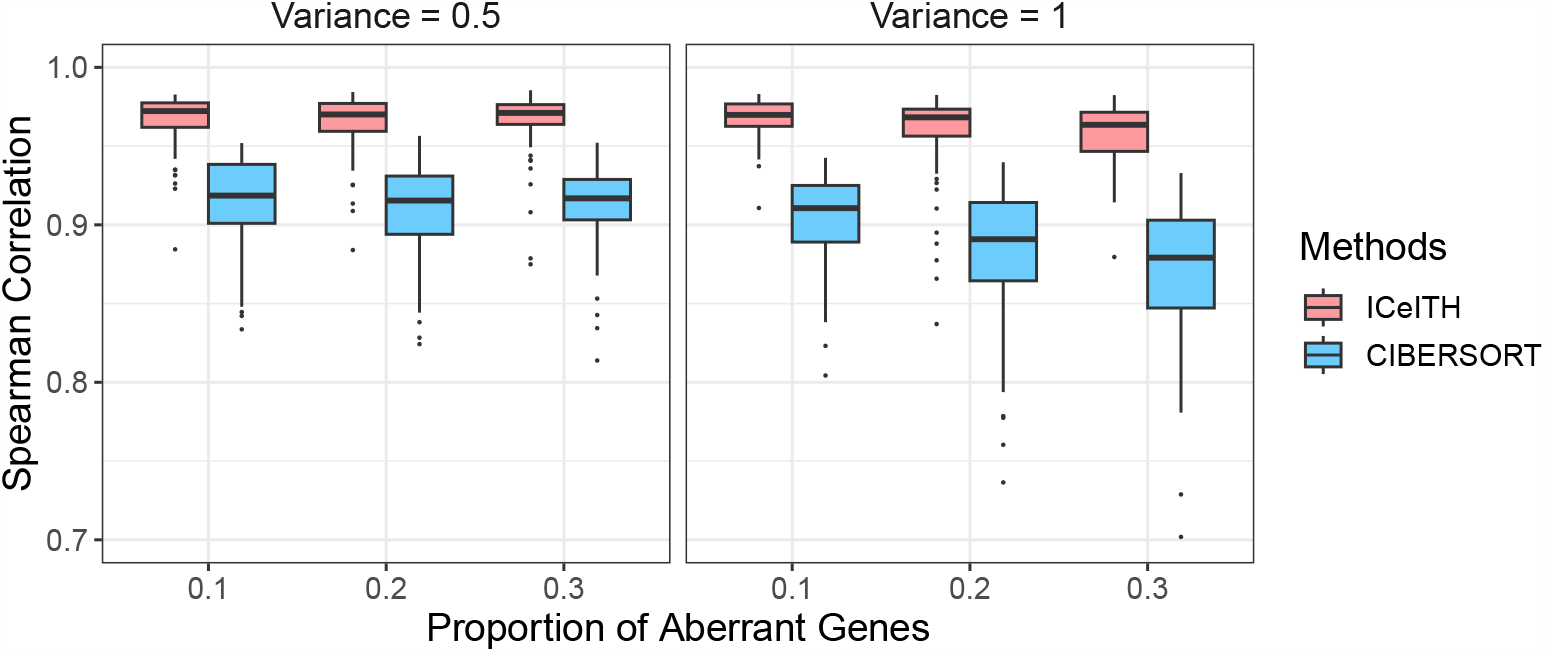
Sampling distribution of Spearman correlation between 4 true CTs (cell types) proportions and estimated CTs from two different methods: ICeITH and CIBERSORT with different proportion/variance of aberrant genes simulated in the reference panel.

In addition, we assessed the performance of our model under varying degrees of within-subject correlation. ICeITH is designed as a hierarchical Bayesian model to address within-subject correlation. From a frequentist standpoint, this is analogous to a linear model with a patient-specific random effect. Consequently, we can express the hidden cell-type-specific gene expression, denoted as *W*_*sgk*_ for sample *s* belonging to patient *i*, as follows: log(*W*_*sgk*_) = *μ*_*gk*_ + *μ*_*igk*_ + *ϵ*_*gk*_. Here, *μ*_*gk*_ represents the mean expression for gene *g* and cell type *k* at a population level, *μ*_*igk*_ is a random effect following a normal distribution with a mean of 0 and variance of 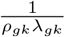 at a patient level, and *ϵ*_*gk*_ represents random noise following another normal distribution with a mean of 0 and variance of 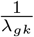.

For any pair of samples *u* and *v* from subject *i*, the covariance between *W*_*ugk*_ and *W*_*vgk*_ is calculated as 1*/*(*ρ*_*gk*_*λ*_*gk*_), and their correlation *ρ*(*W*_*ugk*_, *W*_*vgk*_) is given by 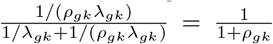. Following the simulation settings outlined in Section 3.1 in the main text, we systematically varied the within-subject correlation across the range of values: 0.01, 0.1, 0.3, 0.5, 0.7, 0.9 and examined the performance of ICeITH under these different conditions.

Figure S4 displays the sampling distribution of Spearman correlations between two methods, ICeITH and CIBERSORT, across various levels of within-subject correlation. The results indicate ICeITH consistently outperforms CIBERSORT, demonstrating its robustness even in the presence of varying degrees of within-subject correlation.

**Figure S4:**
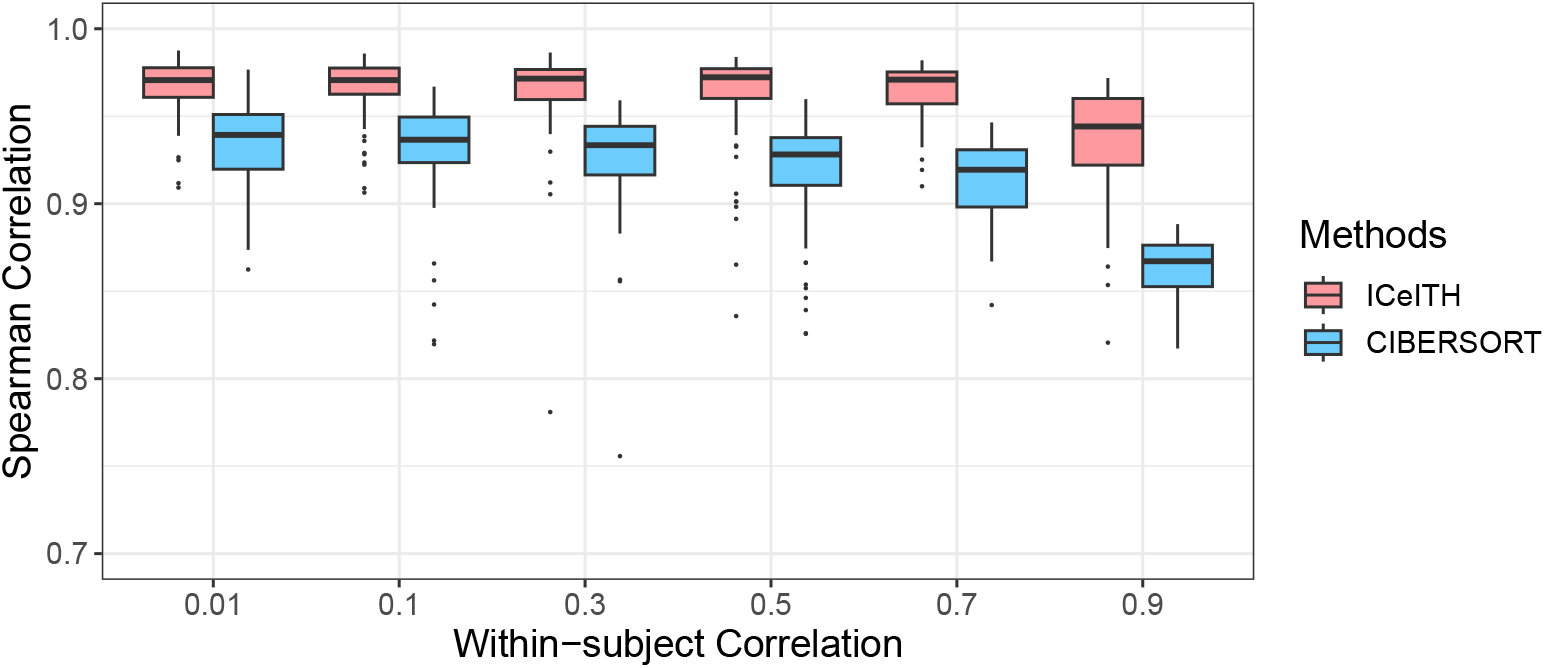
Sampling distribution of Spearman correlation between 4 true CTs (cell types) proportions and estimated CTs from two different methods: ICeITH and CIBERSORT with different degree of within-subject correlation.

## S4 Additional results on real data

### S4.1 Additional material on TRACERx data

**Table S1:** Number of tumor samples per patient from TRACERx cohort, and it only includes patients with more than one RNA-seq data available.

| Number of tumors per patient |  |  |  |  |  |
| --- | --- | --- | --- | --- | --- |
| Number of tumors per patient | 2 | 3 | 4 | 5 | 6 |
| Number of patients | 17 | 14 | 8 | 4 | 2 |

**Figure S5:**
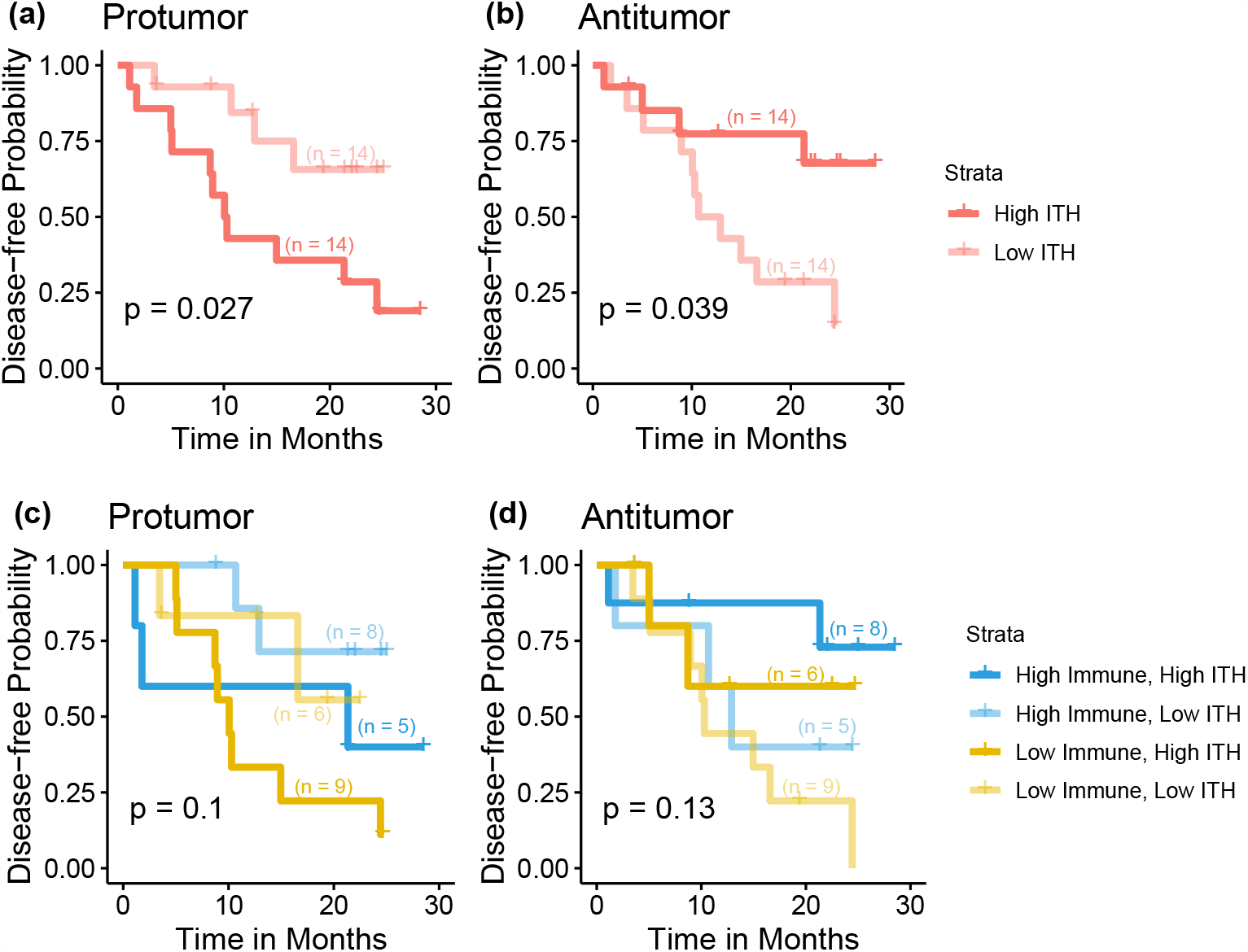
KM (Kaplan-Meier) curves for disease-free probability with high vs. low ITH estimated under pro-(a) and anti-tumor (b) process immune cells for 28 patients from TRACERx study. KM plots for disease-free probability with high vs. low ITH estimated under pro-(c) and anti-tumor (d) process immune cell, stratified by different immune levels, for 28 patients in TRACERx study. The sample size for each stratification is marked on the KM curve.

Motivated by the simulation results that the performance of the ICeITH model increases as more samples are sequenced from each patient, we keep patients (*n* = 28) with more than two regions available and explore the association of intratumor heterogeneity of targeted immune cell types to disease-free survival outcomes. Specifically, for immune cells (i.e., neutrophils and monocytes) that have pro-tumor functions, patients with a higher ITH are at a significantly increased risk compared with a lower ITH (Figure S5(a), *P* value = 0.03). Data from patients stratified by both immune context and ITH further confirms that a higher ITH may be associated with a worse outcome (Figure S5(c)). However, possibly due to the limited sample size, such an effect is not statistically significant (*P* value = 0.1). Meanwhile, patients with high ITH in anti-tumor immune cells (i.e., B cells, CD4T, CD8T and natural killer cells) are at a reduced risk of survival outcome (Figure S5(b), *P* value = 0.04). When we further stratify patients with overall immune context, patients with high ITH still consistently have better survival outcomes (Figure S5(d)). Subsetting patients with a larger sample size provides additional evidence that our proposed method is able to capture the targeted immune cell variability and thus predict prognosis in early-stage lung cancer.

In addition, we have investigated the risk stratification of the TRACERx cohort based on the prevalence of immune cells which are classified as having pro-tumor functions. Specifically, conditional on the protumor immune cell type (e.g. Neutrophils or Monocytes), we measure the immune cell prevalence at the patient level by various summary statistics (e.g. 1st and 3rd Quartile, median, mean, and maximum). Then we dichotomize patients into different risk levels by the median values of these summary statistics and evaluate their association with the disease-free probability as displayed in Figure S6. In general, the different prevalence of Neutrophils cells did not have a strong prognostic association with patients’ survival. However, a higher heterogeneity of Monocytes cells is associated with a worse survival outcome.

**Figure S6:**
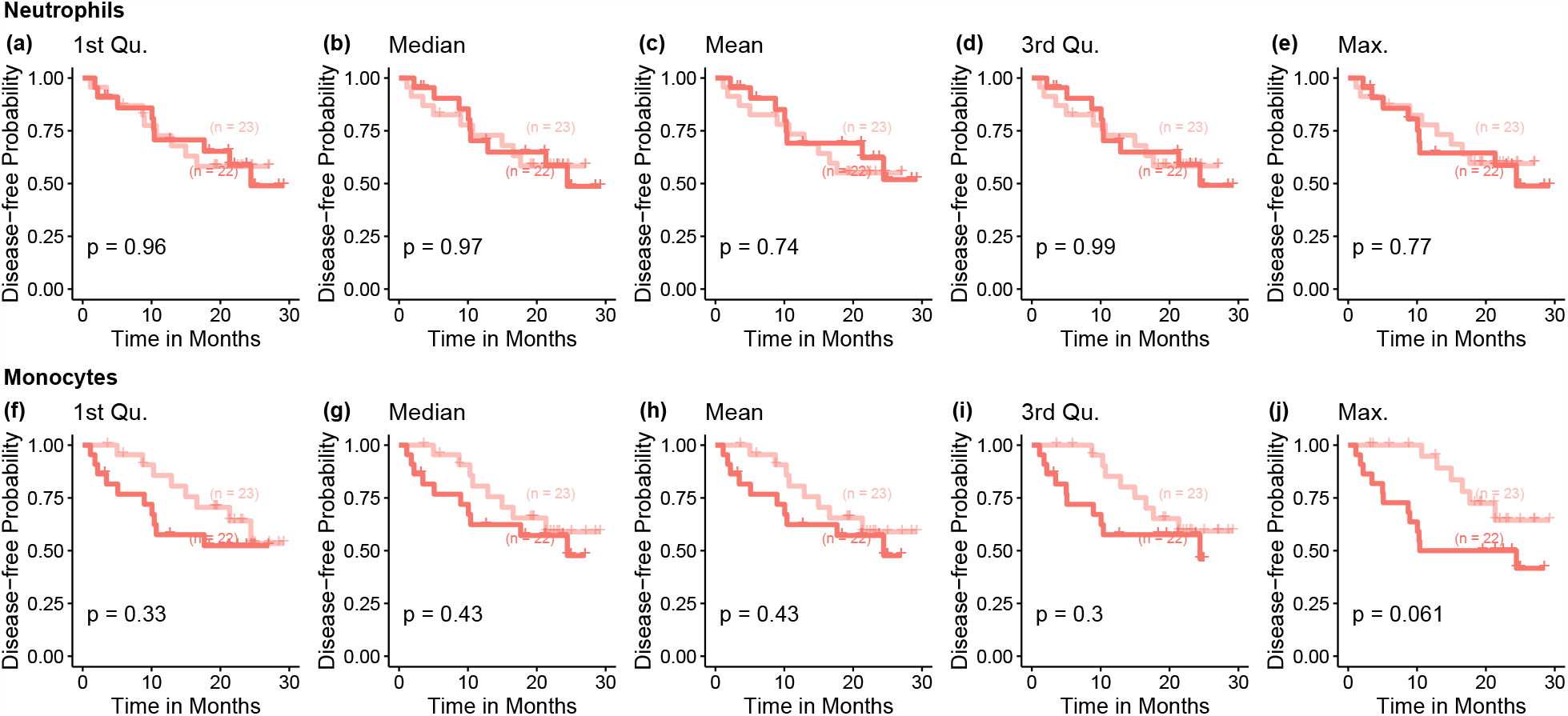
Kaplan-Meier (KM) curves for disease-free probability with high vs. low pro-tumor immune cells prevalence.

### S4.2 Additional material on MDA-ITH data

**Table S2:**
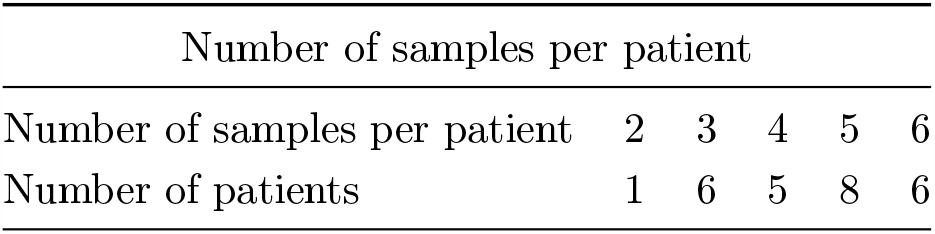
Number of tumor samples per patient from MDA-ITH cohort, and it only includes patients with more than one RNA-seq data available.

